# Auditing Confidence-Calibrated Benchmarking of Single-Cell Perturbation Models

**DOI:** 10.64898/2026.10.12.758706

**Authors:** Manuel Gallone, Shyam Ravidath, Srihith Madireddy

**Author notes:** Equal contribution (co-first authors) (Manuel Gallone, Shyam Ravidath).

## Abstract

A benchmark that reports only the coverage of uncertainty intervals can order uncertainty methods backwards. For scVI on unseen Norman perturbations, MC dropout is the worst-calibrated of three uncertainty quantification (UQ) methods, covering 0.0191 of (cell, gene) pairs at a nominal 0.90, and it is also the best ranking signal: selecting the ten cells it is most confident about reduces DE-gene error by 63.6% against random selection. Calibration and ranking are separate properties and have to be measured separately. We found this while auditing an evaluation harness we had built ourselves, covering two datasets (Norman 2019 and Replogle 2022) in six dataset-split combinations, three models (scVI, CPA and a linear Ridge baseline) and three UQ methods (MC dropout, quantile regression and split conformal prediction), against null baselines the harness had never included. The harness gave each test cell to the model as input and then scored the model on reproducing that same cell. An identity predictor that returns its input unchanged therefore reaches a perturbation-effect correlation of 1.000 on all six splits, so scVI’s 0.27–0.74 is measured against an echo ceiling rather than a null. Against a constant train-mean predictor, scVI wins on 99.7% of Norman cells but only 0.2% of Replogle cells. Two gaps in the harness are stated here rather than later. First, the Replogle split labelled “unseen perturbation” held out sequencing batches, and 2,004 of its 2,007 perturbation labels appear in training. We relabel it “unseen batch” and add a genuine gene holdout, on which every predictor that is denied the target cell is matched or beaten by a constant train-mean profile. Second, the original harness kept no per-cell CPA predictions; regenerated from the saved checkpoints, CPA’s counterfactual prediction, the only deep-model output in the harness that does not read the target cell, has higher error than a constant expression profile on all six splits. Split-conformal coverage held at the nominal 0.900 marginally while covering only 0.271–0.342 of DE-gene intervals, and it fell to 0.847 when the cell line changed from K562 to RPE1. Calibrating a separate quantile per gene set restores in-distribution DE-gene coverage to 0.898–0.901, and recalibrating on RPE1 restores the marginal 0.900. For scVI’s DE genes under that shift both corrections are needed: coverage is 0.766 with neither and 0.901 with both. We document six failure modes, each with its mechanism, the diagnostic that exposes it, and a detection procedure that transfers to other harnesses.

## Introduction

Computational perturbation models predict how cells respond to drugs, gene knockouts, and other interventions, and offer a path toward reducing the experimental burden of drug discovery and functional genomics, a pressing priority given the cost and throughput limits of large-scale wet-lab screens. The data such models learn from are Perturb-seq screens, which pair pooled CRISPR perturbations with a single-cell RNA-seq readout and were first run as CRISPRi screens in K562 cells^1^.

One limitation remains in how these models are assessed. Traditional protocols measure point-prediction accuracy on held-out examples drawn from the training distribution, but perturbation models are most valuable on compounds or cell types not yet experimentally characterized^2^. The meaningful target there is the perturbation effect, the shift in expression relative to matched controls on differentially expressed genes, rather than whole-transcriptome reconstruction, since per-cell all-gene error is dominated by baseline expression and technical variance. Overconfident predictions carry heavy downstream cost, sending resources toward dead ends, so whether a model’s expressed confidence is warranted, formally its calibration, is central to its utility as a decision-support tool. Yet PerturBench^3^ and the broader benchmarking literature remain focused on accuracy; uncertainty is rarely evaluated under distribution shift or tested for decision utility^4^.

We built an OOD evaluation harness to answer which perturbation models generalize most reliably to unseen perturbations and cell types, and how well calibrated their uncertainty estimates are under those conditions. This paper reports what happened when we audited it: against null baselines the harness had never included, most of its headline results turned out to measure something easier than the metric names claim. Beyond calibration error we also evaluate decision utility: given a fixed budget to validate *k* predictions at the bench, does ranking by the model’s own confidence, by conformal or quantile interval width or by predictive variance, beat picking *k* at random? We exclude residual-based confidence on principle, since it requires ground-truth labels and amounts to oracle ranking. We did so while supplying the target cell itself to the models being scored: the same information, entering through a different door^5^. The linear baseline, the only model never given the target cell, is the only one that beats its null on a genuine counterfactual^2^.

### Contributions

The first contribution is that calibration and decision value are dissociated: MC dropout is the worst-calibrated of three UQ methods (0.0191 empirical coverage at a nominal 0.90) and simultaneously the best ranking signal (+63.6% over random selection at *k*=10), so a benchmark that reports only coverage orders these methods backwards. The second is an audit procedure that requires no retraining: we retrofit null baselines to a finished harness, an identity predictor that returns its own input and a constant train-mean predictor that receives no input at all, and every failure mode below was found this way, from stored predictions. The third is six failure modes, each with a mechanism, a diagnostic and a detection procedure that transfers to other harnesses: scoring reconstruction as prediction, metrics a constant already saturates, holding out the wrong column, conditioning on covariates the model never received, silent substitution of unseen categories, and conclusions that depend on a unit convention. The fourth is a measured correction of split conformal rather than a recommendation. Calibrated once on K562 residuals, split-conformal coverage holds marginally in distribution (0.900), covers only 0.271–0.342 of DE-gene intervals, and falls to 0.847 on RPE1. Calibrating a separate quantile per gene set brings DE-gene coverage to 0.898–0.901 in distribution; recalibrating on RPE1 brings marginal coverage back to 0.900; and with both, scVI covers 0.901 of its RPE1 DE-gene intervals (Table 6). The cost is width: on Norman the DE-gene interval is about four times wider than the rest.

### Related work

Perturbation models have evolved from early latent-variable approaches^6^: scGen^7^ showed VAE architectures could generalize across conditions, CPA^8^ extended this to compositional drug-dose-celltype responses, and scVI^9^ introduced a VAE for scRNA-seq conditioned on a batch covariate. All three achieve strong in-distribution performance, but their evaluations emphasize point accuracy rather than calibrated uncertainty. More recent evaluation focuses on the perturbation effect, the correlation of predicted against true expression shifts and the recovery of differentially expressed genes^10^, rather than absolute expression error, which is dominated by the unperturbed baseline. Three independent evaluations on Perturb-seq data of this kind frame our audit: deep perturbation models, among them scGPT, scFoundation, GEARS and CPA, do not yet outperform additive or linear baselines on single and double perturbation effects^2^, a perturbation-informed mean control exceeds GEARS and scGPT^11^, and a training-set-mean baseline beats scGPT and scFoundation^12^.

scPerturb^13^ harmonized 44 public perturbation datasets under a unified QC pipeline, enabling cross-study comparison without preprocessing confounds, but it is a data resource rather than an evaluation framework; it defines no splits or generalization criteria. The closest prior work is PerturBench^3^, which benchmarks multiple models across structured held-out splits but measures only point-prediction quality: it does not test whether uncertainty estimates degrade under shift, or whether confidence scores can prioritize experiments. We treat calibration and decision utility as primary metrics, applying MC dropout^14^, quantile regression^15^ via the pinball loss, and split conformal prediction^16^, following Angelopoulos and Bates ^17^. PertEval-scFM^18^ does test distribution shift, and finds that zero-shot foundation-model embeddings give little gain over simple baselines for perturbation-effect prediction, with the gap widest under shift.

## Methods

Figure 3 (Supporting Online Material) summarises the harness and Table 6 (Supporting Online Material) gives every split. Both datasets were accessed through scPerturb, filtered to drop cells with fewer than 200 detected genes or more than 10% mitochondrial reads, library-size normalised to 1e4, log1p transformed, and reduced to the top 5,000 highly variable genes; on Replogle the HVG set is the intersection common to both cell lines. Norman 2019^19^ (CRISPRa, K562) offers only the in-distribution and unseen-perturbation splits, since it is a single cell line. Replogle 2022^20^ (CRISPRi, K562 and RPE1) supplies the two shift splits. In-distribution scores a random 10% of K562 cells whose perturbations, sequencing batches and cell type all appear in training. Unseen perturbation holds out 20% of guide identities on Norman. The Replogle split configured under that name held out 12 of 48 K562 sequencing batches rather than perturbation identity (“Holding out the wrong column”); we call it *unseen batch* throughout, and add a genuine gene holdout, described below. Unseen cell type scores all RPE1 cells with models trained on K562 alone. Combined OOD scores the RPE1 cells that fall in the held-out batches. The scVI model conditions at inference on the cell-type/batch covariate it was trained on, and no RPE1 cells were used in training, so the unseen-cell-type and combined-OOD splits reflect expression-space shift under a substituted cell-type label rather than held-out covariate conditioning. The linear baseline is given one-hot conditioning on the column each dataset’s split was configured with: perturbation identity on Norman, and sequencing batch on Replogle, where the configuration named batch. It receives no cell-type covariate. All data are public, de-identified single-cell datasets; no human subjects were involved and no IRB review was required.

### A genuine gene holdout on Replogle

To test perturbation novelty on Replogle, we rebuilt the split from the same 80,000 K562 cells, recovering raw counts from the scPerturb source file (the rebuilt log1p matrix matches the stored one to 3.3 × 10^−7^), and held out 20% of the 2,050 gene labels: 410 genes, stratified on whether the gene carries a DE list, with all of their 16,107 cells in the test set. The remaining cells are split 80/10/10 into training, calibration and in-distribution test, and the 2,731 non-targeting control cells are split 70/10/10/10 so that every test set has a control pool. All 410 held-out labels, and every perturbed test cell, are absent from training (Table 10, Supporting Online Material); every split spans all 48 batches. scVI and CPA were retrained on this split, seed 42 only. scVI repeats the original Replogle configuration (128 latent dimensions, two layers, dropout 0.1, batch size 256, at most 200 epochs with early-stopping patience 15, log1p input, cell_type as batch key and no perturbation covariate; scvi-tools 1.5.2), and a second scVI arm adds gene as a categorical covariate and is trained on raw counts. CPA uses the configuration of Table 7 with gene as perturbation key. Neither model stopped early: scVI ran all 200 epochs and CPA all 80. Calibration-split predictions were saved for all three this time. A held-out gene is not in either model’s perturbation vocabulary, so at inference it is mapped explicitly to the non-targeting label, and the number of cells so mapped is logged rather than substituted silently (“Silent substitution of unseen categories”); on the gene-holdout test set that is all 15,819 perturbed cells. Ridge is refitted on a one-hot encoding of gene, the column the split was built on. Results are in Table 3.

### Models

We evaluated three models spanning a range of architectural complexity: scVI^9^ via scvi-tools^21^, CPA^8^ via cpa-tools 0.8.8, and Ridge regression on one-hot conditioning as a lower-bound reference, on perturbation identity for Norman and on sequencing batch for Replogle. Table 7 (Supporting Online Material) gives every training setting. Three choices are arguments rather than settings and belong here. First, scVI predictions come from model.get_normalized_expression(), which returns proportion space; under convention D predictions and targets are mapped symmetrically to log1p using a fixed mean training-library size, 2,273.02 for Norman and 6,342.42 for Replogle, so no re-derived scVI or Ridge metric uses any test-target information. Second, CPA applied a negative-binomial likelihood to log1p-normalised input rather than to raw counts, so our CPA results characterise that configuration rather than the architecture. Third, on Norman CPA was trained on guide_id with every guide pair containing a NegCtrl guide collapsed to a single control label (the 3 control pairs and the 125 single-gene perturbations, 46,248 of 73,200 training cells), so that it learned only the 104 two-gene perturbations as perturbations, whereas on Replogle it received no usable perturbation signal: zeroing the perturbation embedding leaves DE-gene MSE unchanged, and the cited RPL3/KIF11 cosine of −0.115 lies inside the background embedding distribution, where 10.0% of random pairs are at least that separated. Under this configuration the perturbation pathway contributes no measurable advantage; we report it as a negative result and give the numbers in “Conditioning covariates absent at training time”.

CPA’s per-cell predictions were not retained by the original harness, and we regenerated them for this revision from the saved seed-42 checkpoints, in the training environment of Table 7, over every cell of the six evaluation splits. The *reconstruction* arm gives CPA each test cell’s own expression with its perturbation label, the original evaluation protocol; the *counterfactual* arm gives it a control cell drawn at random from the same split, with the target cell’s label, and is the only deep-model prediction in the harness that does not read the target cell. The regeneration matches the earlier audit of the same checkpoint, a Norman in-distribution DE-gene MSE of 0.2634 over 3,385 two-gene-perturbation cells, to six decimals. Labels outside a checkpoint’s vocabulary are fed to CPA as control, explicitly and counted: 8,055 of 19,947 Norman unseen-perturbation cells, which with the NegCtrl collapse leaves no cell of that split with a perturbation label, and 7,942 and 4,366 RPE1 cells on the unseen-cell-type and combined-OOD splits. CPA rows are reported in Tables 2, 3, 5 and 6; the dissociation and decision-value tables and their two figures report scVI alone, because CPA’s uncertainty methods were not rerun. Because CPA applied its negative-binomial likelihood to log1p-normalised input rather than to raw counts, the CPA numbers that do appear characterise that configuration and are not a comparison of the three architectures under identical conditions.

### Uncertainty Quantification and Evaluation

Point and uncertainty estimates are the mean and variance of repeated forward passes; Table 7 gives the pass counts, the quantile-head configuration and the bootstrap settings. For conformal prediction the nonconformity score is the absolute residual of a single (cell, gene) element, so *q*^is one scalar per model and split: the ⌈(*n*+1)(1−*α*)⌉*/n* empirical quantile of the pooled |*y_ij_* − *y*^*_ij_*| over all calibration (cell, gene) pairs. Reported coverage is the marginal fraction of test (cell, gene) pairs inside ± *q*^. The guarantee holds under exchangeability, which is not satisfied when the target cell type is absent from the calibration set. On Replogle the designated calibration split was never scored, since no predictions over those cells were saved, so *q*^ is calibrated on a disjoint random half of the in-distribution test split and evaluated on the other half; the two halves are exchangeable draws from the same K562 pool, with identical cell type, all 36 sequencing batches and no shared cells, and the empirical marginal coverage target is recovered on the held-out half (0.900 [0.899, 0.901]). Because genes within a cell are correlated, the pooled element-level construction does not inherit the finite-sample split-conformal guarantee at the element level; the coverage we report is empirical. To assess decision value we rank predictions by interval width and compute DE-gene MSE over the top *k* against random selection. Both MSE and Pearson *r* are reported over all genes, alongside MSE on the top-20 differentially expressed genes per perturbation; empirical coverage and mean interval width measure uncertainty quality. The DE genes are the 20 gene symbols stored per perturbation in the harness’s DE lists, 197 Norman perturbations in results/metrics/srihith_de_genes_norman.csv and 916 Replogle perturbations in results/replogle_metrics/srihith_de_genes_replogle.csv, and DE-gene MSE is computed per cell over the 20 genes listed for that cell’s perturbation and then averaged over cells, excluding cells whose perturbation has no stored list. The statistic that ranked the genes into those lists is not part of the released re-derivation code, which consumes the lists as stored, and the same lists are used for every DE-gene number in this paper.

The normalized conformal interval that the decision-value rows rank by is a second construction, scored per cell rather than per (cell, gene) element. Its nonconformity score for cell *i* is the mean over the 5,000 genes of |*y_ig_* − *y*^*_ig_*| divided by a difficulty estimate *σ*^(*y*^*_i_*)^22^. *σ*^ is a ridge regression (*α* = 1) of each calibration cell’s mean absolute residual on its 5,000-dimensional prediction vector, standardised per gene and floored at 10*^−^*^3^, fitted on the calibration half of the in-distribution scVI array, 1,000 of the 2,000 Norman cells and 3,025 of the 6,051 Replogle cells, chosen by the same fixed permutation as *q*^. *q*^*_n_* is the ⌈(*n*+1)(1−*α*)⌉*/n* quantile of the calibration scores (1.0018 on Norman, 1.0539 on Replogle), the half-width for a test cell is *q*^*_n_σ*^(*y*^*_i_*), and the ranking signal is 2*q*^*_n_σ*^(*y*^*_i_*). At test time *σ*^ takes the prediction and nothing else, and under convention D the prediction is log(1 + *p_i_L̄*) with *p_i_* the model’s proportion-space output and *L̄* the training-split mean library, so neither the label nor the test cell’s library enters the width. Plain split conformal yields one *q*^, and therefore one width 2*q*^, per model and split by construction, so it cannot rank cells; the normalized interval yields a per-cell width. The in-distribution normalized-conformal rows of Table 7 rank only the evaluation half of the split, 909 DE-evaluable Norman cells and 2,291 Replogle cells, excluding the calibration cells the difficulty model was fitted on; the original-heads rows of the same splits still rank every DE-evaluable cell, 1,808 and 4,615. Ranking the calibration cells with a difficulty model fitted on them would let evaluation data enter model fitting, a form of leakage in the taxonomy of Kapoor and Narayanan^5^, which is why they are excluded.

Two sets of quantile heads appear in this paper. The original heads are three unconstrained linear heads on scVI’s latent *z*, trained for 30 epochs (Adam, learning rate 10^−3^, batch 1,024) on 15,000 training cells at a nominal level of 0.80 (quantiles 0.10, 0.50, 0.90); their stored per-cell widths, the mean over genes of *q*_0.90_ − *q*_0.10_, are negative for 58 of the 2,000 Norman unseen-perturbation cells, and they feed the rows labelled “original heads” in Tables 4 and 7. The refit heads take the top-10 principal components of scVI’s predicted profile under convention D as features, are fitted on the fitting half of the matched in-distribution cells at a nominal level of 0.90 (quantiles 0.05, 0.50, 0.95) with 300 Adam steps from a warm start at the empirical per-gene quantiles, and are made monotone by sorting the three predicted surfaces elementwise at prediction time^23^; they feed the rows labelled “refit heads” in the same two tables.

A per-cell win against the train-mean floor is a cell whose all-gene MSE under the model is strictly smaller than under the constant train-mean profile, ties counting as non-wins; the 99.7% of the abstract is 1,994 wins of 2,000 Norman in-distribution scVI cells and the 0.2% is 10 wins of 6,051 Replogle in-distribution scVI cells.

Confidence intervals on decision value and on conformal coverage resample perturbations with replacement, 2,000 resamples; for Replogle a perturbation-within-batch resampling was also run and is reported in the table footnotes^24^. Significance in Table 7 is by Benjamini–Hochberg^25^ at *q* = 0.05 across the 48 split × signal × *k* comparisons, using the perturbation-cluster bootstrap *p*-values. The Norman unseen-perturbation split has 15 perturbation clusters among its 483 DE-evaluable cells.

### Code and data availability

All datasets are public: Norman 2019 and Replogle 2022, both accessed through scPerturb. The re-derivation code and the per-split result tables behind the tables and figures of the original submission are available at https://anonymous.4open.science/r/audit-scmodel-paper-AF8D/, and each of them regenerates from those result tables alone, with no model and no GPU. The scripts and result tables added in this revision (the CPA rows, the gene-holdout analysis, Table 5 and Table 11) are being added to the same release and are available from the authors until then. Per-cell prediction arrays (9.2 GB) and model checkpoints are too large to host and are available from the authors on request. Table 1 lists, for every table and figure, the released result file it is typeset from and what recomputing that file requires.

**Table 1:**
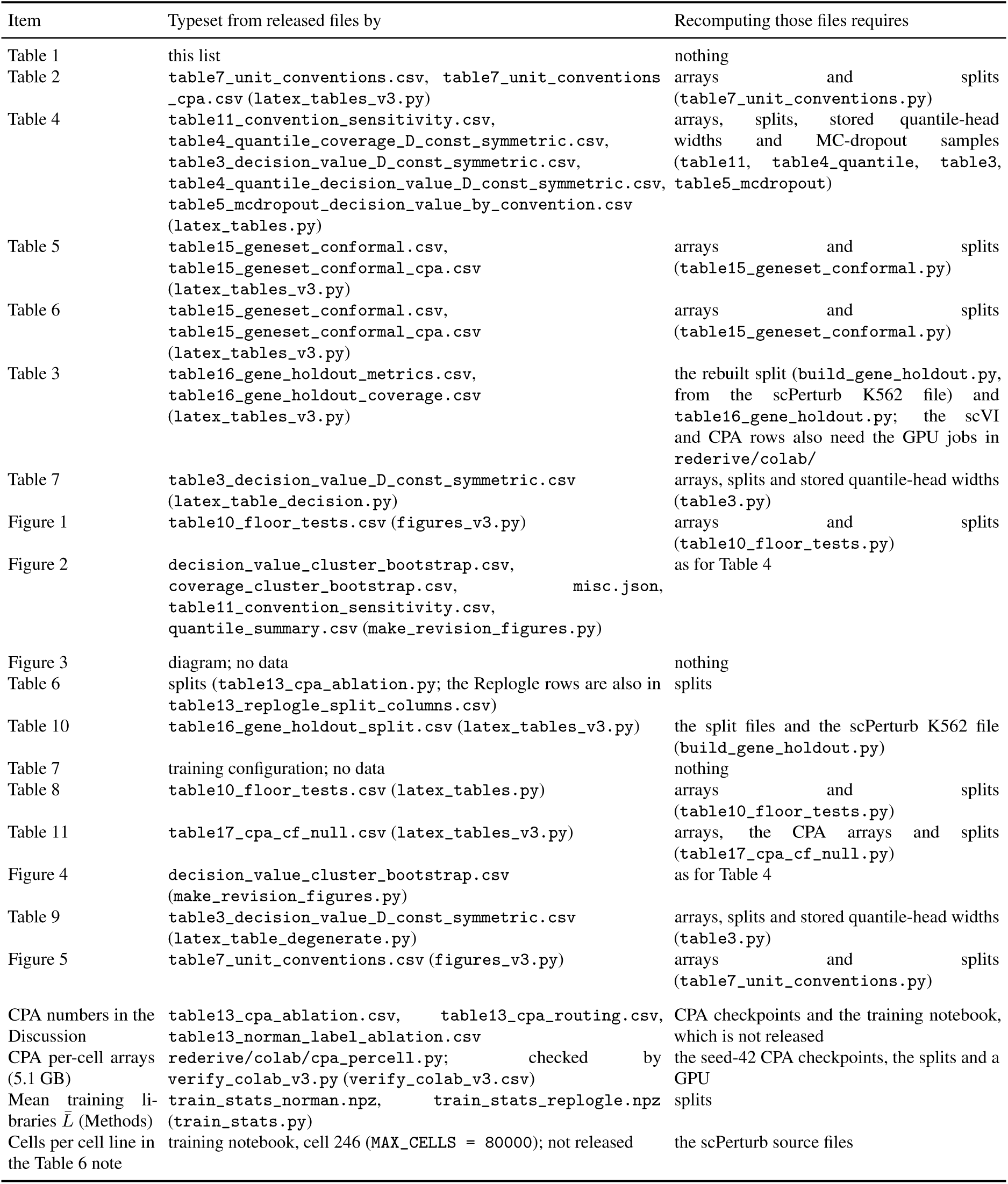
What each table and figure regenerates from. Released result files are the CSVs in results_rederived_v2/ of the code repository; “arrays” are the per-cell prediction arrays (9.2 GB) and “splits” the split .h5ad files, both available on request rather than released. The perturbation-level intervals, Benjamini–Hochberg adjusted *p*-values, normalized-interval coverage and plain-conformal width added in revision are computed from the arrays and splits by the revision audit script reviewer_audit.py, whose outputs accompany this revision.

| Item | Typeset from released files by | Recomputing those files requires |
|---|---|---|
| Table 1 | this list | nothing |
| Table 2 | table7_unit_conventions.csv, table7_unit_conventions_cpa.csv (latex_tables_v3.py) | arrays and splits (table7_unit_conventions.py) |
| Table 4 | table11_convention_sensitivity.csv, table4_quantile_coverage_D_const_symmetric.csv, table3_decision_value_D_const_symmetric.csv, table4_quantile_decision_value_D_const_symmetric.csv, table5_mcdropout_decision_value_by_convention.csv (latex_tables.py) | arrays, splits, stored quantile-head widths and MC-dropout samples (table11, table4_quantile, table3, table5_mcdropout) |
| Table 5 | table15_geneset_conformal.csv, table15_geneset_conformal_cpa.csv (latex_tables_v3.py) | arrays and splits (table15_geneset_conformal.py) |
| Table 6 | table15_geneset_conformal.csv, table15_geneset_conformal_cpa.csv (latex_tables_v3.py) | arrays and splits (table15_geneset_conformal.py) |
| Table 3 | table16_gene_holdout_metrics.csv, table16_gene_holdout_coverage.csv (latex_tables_v3.py) | the rebuilt split (build_gene_holdout.py, from the scPerturb K562 file) and table16_gene_holdout.py; the scVI and CPA rows also need the GPU jobs in rederive/colab/ |
| Table 7 | table3_decision_value_D_const_symmetric.csv (latex_table_decision.py) | arrays, splits and stored quantile-head widths (table3.py) |
| Figure 1 | table10_floor_tests.csv (figures_v3.py) | arrays and splits (table10_floor_tests.py) |
| Figure 2 | decision_value_cluster_bootstrap.csv, coverage_cluster_bootstrap.csv, misc.json, table11_convention_sensitivity.csv, quantile_summary.csv (make_revision_figures.py) | as for Table 4 |
| Figure 3 | diagram; no data | nothing |
| Table 6 | splits (table13_cpa_ablation.py; the Replogle rows are also in table13_replogle_split_columns.csv) | splits |
| Table 10 | table16_gene_holdout_split.csv (latex_tables_v3.py) | the split files and the scPerturb K562 file (build_gene_holdout.py) |
| Table 7 | training configuration; no data | nothing |
| Table 8 | table10_floor_tests.csv (latex_tables.py) | arrays and splits (table10_floor_tests.py) |
| Table 11 | table17_cpa_cf_null.csv (latex_tables_v3.py) | arrays, the CPA arrays and splits (table17_cpa_cf_null.py) |
| Figure 4 | decision_value_cluster_bootstrap.csv (make_revision_figures.py) | as for Table 4 |
| Table 9 | table3_decision_value_D_const_symmetric.csv (latex_table_degenerate.py) | arrays, splits and stored quantile-head widths (table3.py) |
| Figure 5 | table7_unit_conventions.csv (figures_v3.py) | arrays and splits (table7_unit_conventions.py) |
| CPA numbers in the Discussion | table13_cpa_ablation.csv, table13_cpa_routing.csv, table13_norman_label_ablation.csv | CPA checkpoints and the training notebook, which is not released |
| CPA per-cell arrays (5.1 GB) | rederive/colab/cpa_percell.py; checked by verify_colab_v3.py (verify_colab_v3.csv) | the seed-42 CPA checkpoints, the splits and a GPU |
| Mean training li-braries *L̄* (Methods) | train_stats_norman.npz, train_stats_replogle.npz (train_stats.py) | splits |
| Cells per cell line in the Table 6 note | training notebook, cell 246 (MAX_CELLS = 80000); not released | the scPerturb source files |

## Results

### Predictive performance and perturbation effect

We evaluated scVI, CPA and the linear baseline on Norman 2019 and Replogle 2022 across four splits and three UQ methods; Table 2 reports all results. DE-gene MSE on the top-20 differentially expressed genes per perturbation is the primary accuracy metric, since those genes are the most biologically affected and their large expression changes dominate all-gene MSE. scVI and the linear baseline are close on all-gene MSE and Pearson *r* on both Norman splits (0.0226 against 0.0277 and 0.9289 against 0.9119 in distribution; 0.0233 against 0.0294 and 0.9267 against 0.9060 under unseen perturbation) but not on DE-gene MSE, where Ridge scores 0.3291 against scVI’s 0.1921 in distribution and 0.4056 against 0.2148 under unseen perturbation. CPA’s two arms bracket the question the identity row raises. Its reconstruction, which reads the target cell, behaves like scVI on Norman (DE-gene MSE 0.2478 against scVI’s 0.1921 in distribution). On Replogle, where scVI loses to the floor on all-gene MSE on every split, the reconstruction beats the floor on both all-gene and DE-gene MSE on all four (0.2635 against 0.3386 DE-gene MSE in distribution); both are read against the echo ceiling. Its counterfactual, which does not read the target cell, has a higher all-gene and DE-gene MSE than the train-mean floor on all four K562 rows (on Norman in distribution, DE-gene MSE 0.5332 against 0.4964), and carries perturbation signal only on Norman in distribution (effect *r* = 0.188; −0.079 to 0.025 elsewhere). On the two RPE1 rows it does beat the train-mean floor, but only because its input cells are RPE1 controls: the mean of those same control cells, a constant, has a DE-gene MSE of 0.2011 and 0.1993 against the counterfactual’s 0.3133 and 0.3117 (Table 11, Supporting Online Material). Once denied the target cell, CPA beats a constant on no split; on Replogle this is what a checkpoint whose perturbation pathway confers no measurable advantage would produce (“Models”). On Replogle, scVI and Ridge both degrade under OOD conditions, and unseen-celltype and combined-OOD are near-identical, which identifies cell-type shift rather than perturbation novelty as the driver of the loss^18^.

**Table 2:** Point accuracy under the constant-library convention (D), which gives no predictor any test-target information, bracketed by a null and a ceiling. The train-mean floor is a constant profile computed on the training split; it receives no input at all ^11,12^. The identity predictor returns each test cell’s own measured expression unchanged. It is the ceiling for any model whose encoder reads the target cell, which every scVI row and every CPA reconstruction row here does, so those rows are read against an echo ceiling rather than a null. The CPA counterfactual is the only deep-model row that does not read the target cell: its input is a control cell drawn at random from the same split, with the target cell’s perturbation label ^8^. Pearson *r* is the flattened element-wise correlation over all (cell, gene) pairs, the statistic the original study reported. Effect *r* is, for each perturbation, the correlation over its 20 DE genes of the predicted effect (predicted perturbation mean minus predicted control mean) with the observed effect, averaged over perturbations. All-gene MSE is over all (cell, gene) elements of the split; DE-gene MSE is computed per cell over the 20 DE genes of that cell’s perturbation and averaged over cells, and its CI is a bootstrap over cells.

| Split | Predictor | all-gene MSE | Pearson *r* | DE-gene MSE | 95% CI | effect *r* |
|---|---|---|---|---|---|---|
| Norman, in-distribution | scVI | 0.0226 | 0.9289 | 0.1921 | [0.1870, 0.1978] | 0.7437 |
|  | CPA, reconstruction | 0.0239 | 0.9251 | 0.2478 | [0.2378, 0.2592] | 0.6423 |
|  | CPA, counterfactual | 0.0349 | 0.8912 | 0.5332 | [0.5106, 0.5559] | 0.1875 |
|  | Ridge | 0.0277 | 0.9119 | 0.3291 | [0.3182, 0.3396] | 0.6689 |
|  | *train-mean floor* | 0.0297 | 0.9054 | 0.4964 | [0.4738, 0.5173] | – |
|  | *identity (echo ceiling)* | 0.0004 | 0.9987 | 0.0074 | [0.0064, 0.0085] | 0.9978 |
| Norman, unseen perturbation | scVI | 0.0233 | 0.9267 | 0.2148 | [0.2029, 0.2274] | 0.6314 |
|  | CPA, reconstruction | 0.0237 | 0.9255 | 0.2212 | [0.2090, 0.2333] | 0.6728 |
|  | CPA, counterfactual | 0.0354 | 0.8902 | 0.4669 | [0.4254, 0.5098] | -0.0792 |
|  | Ridge | 0.0294 | 0.9060 | 0.4056 | [0.3691, 0.4429] | – |
|  | *train-mean floor* | 0.0294 | 0.9061 | 0.4107 | [0.3734, 0.4493] | – |
|  | *identity (echo ceiling)* | 0.0004 | 0.9988 | 0.0069 | [0.0053, 0.0087] | 0.9968 |
| Replogle, in-distribution | scVI | 0.2409 | 0.7578 | 0.3288 | [0.3198, 0.3380] | 0.2890 |
|  | CPA, reconstruction | 0.1356 | 0.8185 | 0.2635 | [0.2591, 0.2681] | 0.2760 |
|  | CPA, counterfactual | 0.1582 | 0.7862 | 0.3611 | [0.3543, 0.3674] | -0.0108 |
|  | Ridge | 0.1456 | 0.8030 | 0.3353 | [0.3294, 0.3412] | 0.0018 |
|  | *train-mean floor* | 0.1470 | 0.8010 | 0.3386 | [0.3328, 0.3446] | – |
|  | *identity (echo ceiling)* | 0.0001 | 0.9999 | 0.0004 | [0.0004, 0.0004] | 0.9998 |
| Replogle, unseen batch | scVI | 0.2392 | 0.7573 | 0.4203 | [0.4152, 0.4253] | 0.3070 |
|  | CPA, reconstruction | 0.1368 | 0.8167 | 0.1711 | [0.1693, 0.1729] | 0.3877 |
|  | CPA, counterfactual | 0.1585 | 0.7855 | 0.2261 | [0.2230, 0.2291] | 0.0249 |
|  | Ridge | 0.1475 | 0.7995 | 0.2065 | [0.2038, 0.2093] | – |
|  | *train-mean floor* | 0.1475 | 0.7996 | 0.2065 | [0.2038, 0.2093] | – |
|  | *identity (echo ceiling)* | 0.0001 | 0.9999 | 0.0003 | [0.0003, 0.0003] | 0.9998 |
| Replogle, unseen cell type | scVI | 0.3242 | 0.6558 | 0.4761 | [0.4728, 0.4797] | 0.2880 |
|  | CPA, reconstruction | 0.2193 | 0.7185 | 0.2776 | [0.2757, 0.2796] | 0.3371 |
|  | CPA, counterfactual | 0.2444 | 0.6835 | 0.3133 | [0.3111, 0.3158] | 0.0025 |
|  | Ridge | 0.2518 | 0.6741 | 0.3334 | [0.3309, 0.3366] | 0.0290 |
|  | *train-mean floor* | 0.2522 | 0.6734 | 0.3333 | [0.3308, 0.3365] | – |
|  | *identity (echo ceiling)* | 0.0006 | 0.9998 | 0.0009 | [0.0009, 0.0009] | 0.9997 |
| Replogle, combined OOD | scVI | 0.3191 | 0.6582 | 0.4721 | [0.4674, 0.4767] | 0.2715 |
|  | CPA, reconstruction | 0.2159 | 0.7209 | 0.2744 | [0.2717, 0.2772] | 0.3160 |
|  | CPA, counterfactual | 0.2410 | 0.6856 | 0.3117 | [0.3083, 0.3152] | -0.0190 |
|  | Ridge | 0.2480 | 0.6763 | 0.3301 | [0.3260, 0.3340] | – |
|  | *train-mean floor* | 0.2481 | 0.6762 | 0.3301 | [0.3261, 0.3340] | – |
|  | *identity (echo ceiling)* | 0.0006 | 0.9998 | 0.0010 | [0.0010, 0.0010] | 0.9997 |

Norman rows are the 2,000 cells for which scVI predictions were saved; Replogle rows cover every cell. CPA was rerun from the saved seed-42 checkpoints over every cell and is scored here on the same cells; labels outside a checkpoint's vocabulary are fed to it as control, explicitly (“Silent substitution of unseen categories”). Unconverted, the identity predictor scores exactly 0 MSE and *r* = 1.000 on every split; under convention D its input is rescaled to the training-mean library like every other predictor, which is why its effect *r* is 0.997–1.000 rather than exactly 1. “– marks an effect correlation that is undefined because the prediction has no perturbation-specific variance: the floor by construction, and Ridge wherever its one-hot feature (guide on Norman, sequencing batch on Replogle) is absent from the training vocabulary, so it returns its intercept. The Replogle unseen-batch split holds out 12 of 48 K562 sequencing batches; 2,004 of its 2,007 perturbation labels appear in training (“Holding out the wrong column”).

Table 2 brackets every model between two references. The train-mean floor receives no input at all. The identity predictor receives the test cell and returns it unchanged, which is exactly the information the harness gave scVI and gives CPA’s reconstruction arm. Unconverted, it scores zero error and an effect correlation of 1.000 on every split. Rescaled to the training-mean library under convention D, like every other predictor, it scores 0.997 to 1.000. Against that ceiling, scVI’s effect correlations of 0.27–0.74 measure how much of its own input the model fails to reconstruct, not how much perturbation biology it has learned.

Instead of measuring only raw all-gene expression, we also measured the perturbation effect, the change in expression relative to control on DE genes, since baseline expression overshadows raw MSE and obscures whether the model captured the perturbation. On Norman in-distribution both models predict the effect (scVI 0.744, Ridge 0.865). On unseen perturbation the Ridge effect correlation is undefined rather than small: its one-hot feature row is built against the training vocabulary, so an unseen guide yields an all-zero row and the model returns its intercept, giving a prediction with zero across-cell variance and therefore no effect to correlate. Against a constant train-mean predictor on DE genes (Table 2; the all-gene view is Figure 1 and Table 8, Supporting Online Material), scVI improves by 61.3% [58.3, 64.1] in distribution and 47.7% [33.1, 59.1] under unseen perturbation, the latter over 483 measurable cells spanning 15 perturbations, since only 15 of the 58 held-out guides carry a differential-expression list.

**Figure 1:**
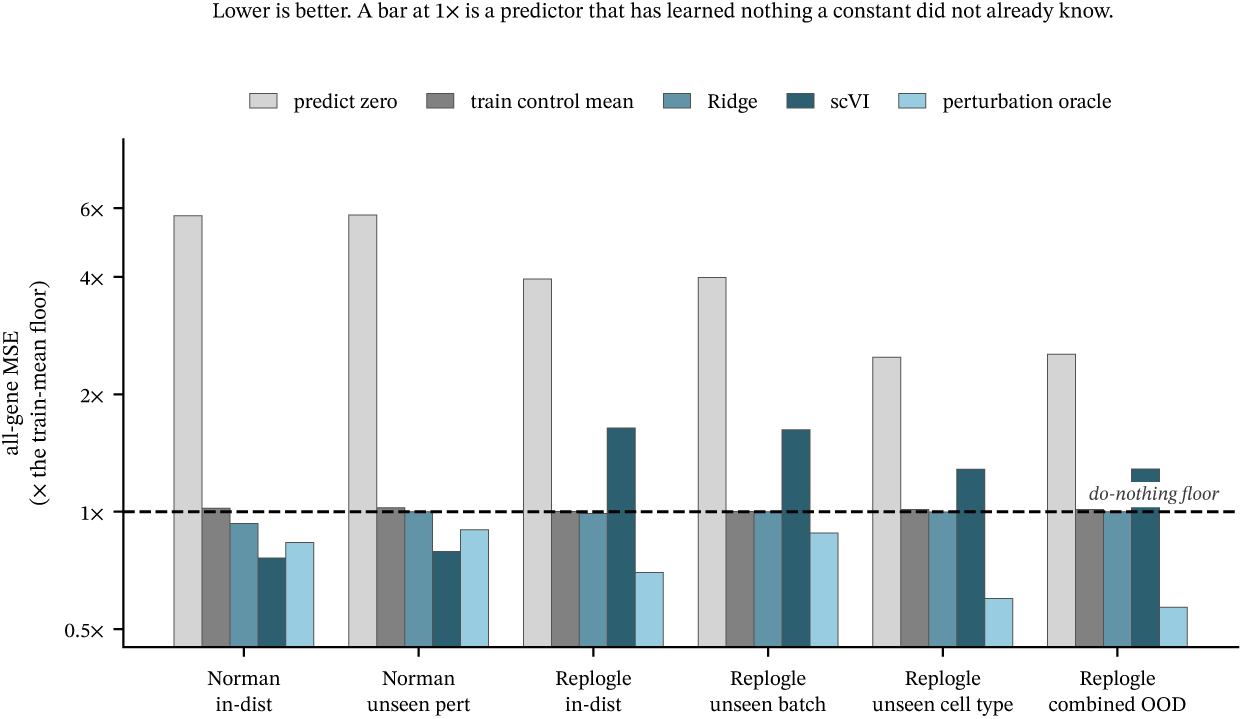
Every predictor as a multiple of the constant train-mean floor, under convention D. Lower is better; a bar at 1× is a predictor that has learned nothing a constant did not already know. scVI clears the floor on both Norman splits and sits above it on all four Replogle splits, where neither model receives perturbation identity. Plotted with matplotlib. Each bar is the mean per-cell all-gene MSE of a predictor divided by that of the train-mean floor.

On the rebuilt Replogle gene holdout (Table 3), Ridge conditioned on gene, the column the split was built on, carries perturbation signal in distribution, which it never did when conditioned on sequencing batch. Its DE-gene MSE is 0.2133 against the floor’s 0.2201, and its effect correlation is 0.197, against 0.002 under batch conditioning. On the 410 held-out genes it returns its intercept and matches the floor on every metric (0.1461 all-gene MSE for both). That row also shows a limit of effect correlation as a metric. Ridge still scores 0.140 on the held-out genes while knowing nothing about them, because its intercept, the mean of mostly perturbed training cells, differs from its control prediction by a generic perturbation response that correlates with each specific effect. An effect correlation therefore needs a null of its own, the training perturbed-mean minus the control mean, before it can be read as perturbation-specific learning.

**Table 3:**
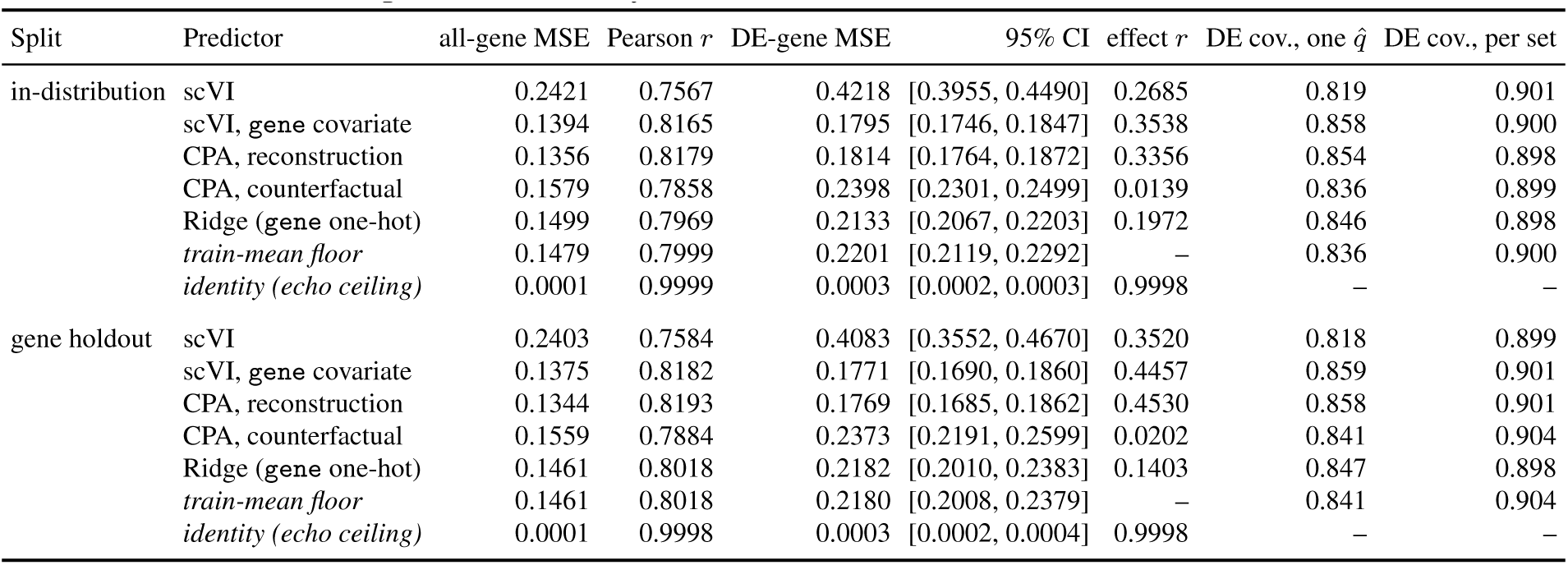
Every predictor on the rebuilt Replogle K562 split (Table 10), under convention D with L̄ recomputed on its own training split. Columns as in Table 2; the DE-gene CI resamples perturbations. The two coverage columns are DE-gene split-conformal coverage at a nominal 0.90, calibrated on this split’s own calibration cells with one *q*^and per gene set (Table 6). Held-out genes are fed to scVI and CPA as the non-targeting label, explicitly; Ridge returns its intercept for them by construction. Both scVI rows and the CPA reconstruction read each test cell’s own expression and are bounded by the identity row; the CPA counterfactual starts from a control cell of the same split. Seed 42 only.

| Split | Predictor | all-gene MSE | Pearson *r* | DE-gene MSE | 95% CI | effect *r* | DE cov., one *q̂* | DE cov., per set |
|---|---|---|---|---|---|---|---|---|
| in-distribution | scVI | 0.2421 | 0.7567 | 0.4218 | [0.3955, 0.4490] | 0.2685 | 0.819 | 0.901 |
|  | scVI, gene covariate | 0.1394 | 0.8165 | 0.1795 | [0.1746, 0.1847] | 0.3538 | 0.858 | 0.900 |
|  | CPA, reconstruction | 0.1356 | 0.8179 | 0.1814 | [0.1764, 0.1872] | 0.3356 | 0.854 | 0.898 |
|  | CPA, counterfactual | 0.1579 | 0.7858 | 0.2398 | [0.2301, 0.2499] | 0.0139 | 0.836 | 0.899 |
|  | Ridge (gene one-hot) | 0.1499 | 0.7969 | 0.2133 | [0.2067, 0.2203] | 0.1972 | 0.846 | 0.898 |
|  | *train-mean floor* | 0.1479 | 0.7999 | 0.2201 | [0.2119, 0.2292] | – | 0.836 | 0.900 |
|  | *identity (echo ceiling)* | 0.0001 | 0.9999 | 0.0003 | [0.0002, 0.0003] | 0.9998 | – | – |
| gene holdout | scVI | 0.2403 | 0.7584 | 0.4083 | [0.3552, 0.4670] | 0.3520 | 0.818 | 0.899 |
|  | scVI, gene covariate | 0.1375 | 0.8182 | 0.1771 | [0.1690, 0.1860] | 0.4457 | 0.859 | 0.901 |
|  | CPA, reconstruction | 0.1344 | 0.8193 | 0.1769 | [0.1685, 0.1862] | 0.4530 | 0.858 | 0.901 |
|  | CPA, counterfactual | 0.1559 | 0.7884 | 0.2373 | [0.2191, 0.2599] | 0.0202 | 0.841 | 0.904 |
|  | Ridge (gene one-hot) | 0.1461 | 0.8018 | 0.2182 | [0.2010, 0.2383] | 0.1403 | 0.847 | 0.898 |
|  | *train-mean floor* | 0.1461 | 0.8018 | 0.2180 | [0.2008, 0.2379] | – | 0.841 | 0.904 |
|  | *identity (echo ceiling)* | 0.0001 | 0.9998 | 0.0003 | [0.0002, 0.0004] | 0.9998 | – | – |

Every scVI row and the CPA reconstruction read the test cell, and they look strongest exactly where they should be weakest. On the held-out genes, CPA’s reconstruction reaches an effect correlation of 0.453 and the gene-covariate scVI 0.446, above their in-distribution 0.336 and 0.354, although every one of the 15,819 perturbed held-out cells was given to both models as non-targeting: the effect they reproduce can only have come from the input cell, which is the echo ceiling at work rather than generalisation. The faithful scVI arm, which like the original Replogle scVI has no perturbation covariate, is worse than the floor on every metric except effect correlation (all-gene MSE 0.2403 against 0.1461).

CPA’s counterfactual, the only deep-model prediction here that does not read the target cell, has an effect correlation of 0.020 on the held-out genes and 0.014 in distribution, and a higher all-gene MSE than the floor on both (0.1559 and 0.1579 against 0.1461 and 0.1479). On a genuine perturbation holdout, then, every predictor in the harness that is denied the target cell is matched (Ridge) or beaten (CPA) by a constant profile. Coverage transfers to the new split: calibrated on its own calibration cells, every predictor holds all-gene coverage at 0.901–0.905 on the held-out genes, DE-gene coverage under one *q*^is 0.818–0.859, and calibrating per gene set restores 0.898–0.904.

### Calibration of uncertainty methods

We tested calibration for three UQ methods (conformal, quantile regression, variance-based) across both models and both datasets. All of our coverage numbers are element-level: each (cell, gene) interval is scored separately, rather than requiring every gene in a cell to be covered at once. Conformal was stable at its 0.90 target across both datasets within cell type. The quantile heads as originally configured do not attain any nominal level: three unconstrained linear heads cross on 48.7% of (cell, gene) pairs on Norman scVI, and their coverage is identical at nominal 0.80 and 0.90 (0.2576 vs 0.2576), which is only possible if the head never learned the quantile level. For scVI at nominal 0.90, refitting with a monotonicity constraint^23^ removes crossing entirely (0.0%) and gives coverage of 0.913–0.985 across splits; Ridge remains under-covered or degenerate on several splits.

The variance-based readings turn on a unit convention. Reported as near-zero (∼ 10*^−^*^8^) across all splits, the MC dropout variances were computed in proportion space and never mapped to the log1p space in which every other quantity here is measured; converted correctly they are roughly four orders of magnitude larger, and MC dropout has ample spread to work with. Two effects had been merged under one explanation. For a separate two-row Norman checkpoint artifact the near-zero variance is real and is not dropout at all: the harness called the deterministic get_normalized_expression() twenty times, so the twenty passes are identical and the reported 1.1 × 10^−16^ is floating-point round-off. That value should not be generalized to every CPA result. Coverage for MC dropout was never computed in the original evaluation; measured properly it is 0.0191 at a nominal 0.90 (Table 4), which makes it the worst-calibrated of the three methods rather than a method with no signal. The linear ensemble’s near-zero spread is a separate and genuine finding: five Ridge fits at *α* ∈ {0.5, 1.0, 2.0, 5.0, 10.0} converge to nearly identical solutions, so inter-member spread has nothing to capture.

**Table 4:**
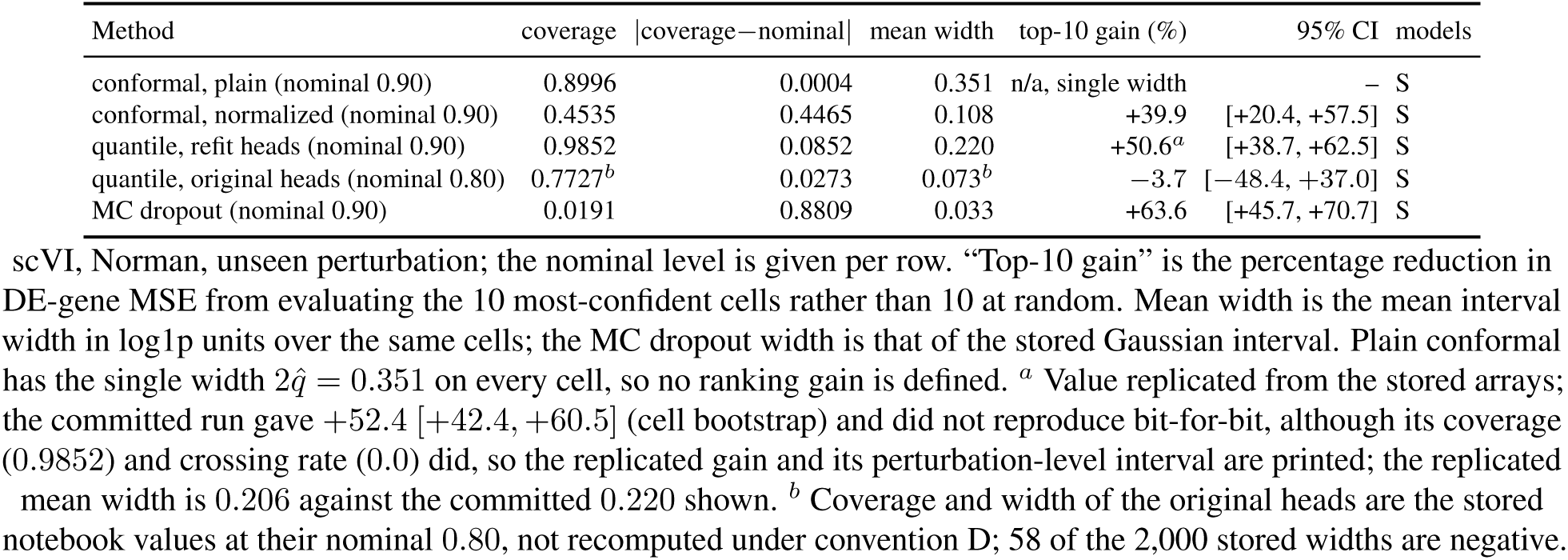
Calibration, interval width and ranking are three separate properties of an uncertainty method, reported side by side: coverage measures calibration, mean interval width measures sharpness, and the top-10 gain measures ranking ^26^. MC dropout is the *worst*-calibrated of the three methods, has the narrowest intervals, and is simultaneously the *best* ranking signal, so no single column orders the methods. Coverage is the fraction of (cell, gene) pairs whose interval covers the truth; the top-10 gain is computed from per-cell DE-gene MSE, so the unit ranked and selected is the cell. For the normalized conformal row, coverage is the fraction of cells whose mean absolute residual lies within *q*^*_n_σ*^, the per-cell quantity that interval is built on. The CI resamples the 15 perturbations present among the 483 DE-evaluable cells, 2,000 resamples. The models column lists the models each row covers (S, scVI; R, Ridge). CPA is in no row: its per-cell predictions were regenerated for this revision (Table 2), but the uncertainty methods scored here were not rerun for it.

| Method | coverage | \|coverage−nominal\| | mean width | top-10 gain (%) | 95% CI | models |
|---|---|---|---|---|---|---|
| conformal, plain (nominal 0.90) | 0.8996 | 0.0004 | 0.351 | n/a, single width | – | S |
| conformal, normalized (nominal 0.90) | 0.4535 | 0.4465 | 0.108 | +39.9 | [+20.4, +57.5] | S |
| quantile, refit heads (nominal 0.90) | 0.9852 | 0.0852 | 0.220 | +50.6<sup>a</sup> | [+38.7, +62.5] | S |
| quantile, original heads (nominal 0.80) | 0.7727<sup>b</sup> | 0.0273 | 0.073<sup>b</sup> | −3.7 | [−48.4, +37.0] | S |
| MC dropout (nominal 0.90) | 0.0191 | 0.8809 | 0.033 | +63.6 | [+45.7, +70.7] | S |

scVI, Norman, unseen perturbation; the nominal level is given per row. “Top-10 gain” is the percentage reduction in DE-gene MSE from evaluating the 10 most-confident cells rather than 10 at random. Mean width is the mean interval width in log1p units over the same cells; the MC dropout width is that of the stored Gaussian interval. Plain conformal has the single width 2q̂ = 0.351 on every cell, so no ranking gain is defined. <sup>a</sup> Value replicated from the stored arrays; the committed run gave +52.4 [+42.4, +60.5] (cell bootstrap) and did not reproduce bit-for-bit, although its coverage (0.9852) and crossing rate (0.0) did, so the replicated gain and its perturbation-level interval are printed; the replicated mean width is 0.206 against the committed 0.220 shown. <sup>b</sup> Coverage and width of the original heads are the stored notebook values at their nominal 0.80, not recomputed under convention D; 58 of the 2,000 stored widths are negative.

### Coverage under distribution shift

Conformal coverage does not hold under cell-type shift (Table 5). Loss of performance under distribution shift is not specific to this harness: PertEval-scFM reports its widest gap between foundation-model embeddings and simple baselines under shift^18^; here the loss appears in coverage. Both models lose coverage on the unseen-cell-type split: scVI falls to 0.847 [0.8464, 0.8476] and Ridge to 0.822 [0.8211, 0.8228], against 0.900 in distribution. This is what conformal theory predicts rather than a defect: the guarantee holds only when calibration and test data are exchangeable, and cell-type shift breaks that^27^. We also measured coverage restricted to the DE genes, using the same *q*^and the same calibrate-evaluate protocol, so both columns come from one computation. On Replogle, scVI covers 0.766 [0.765, 0.768] of (cell, DE-gene) pairs under cell-type shift against 0.847 marginally, so the genes the analysis is about lose coverage faster than the transcriptome as a whole. This DE-gene coverage is an empirical fraction over correlated (cell, DE-gene) pairs and carries no finite-sample guarantee. On Norman the divergence is severe in distribution: both models hold marginal coverage at the nominal 0.900 while covering 0.342 [0.334, 0.351] (scVI) and 0.271 [0.267, 0.275] (Ridge) of DE-gene pairs. These two DE-gene coverages are likewise empirical fractions over correlated (cell, DE-gene) pairs with no finite-sample guarantee, and the Norman scVI and Ridge rows are not scored on the same cells. A marginal guarantee at 0.90 is compatible with covering roughly a quarter to a third of the intervals a prioritization decision rests on, and split conformal promises nothing else: the DE genes are exactly the high-residual subset a marginal quantile averages away.

**Table 5:**
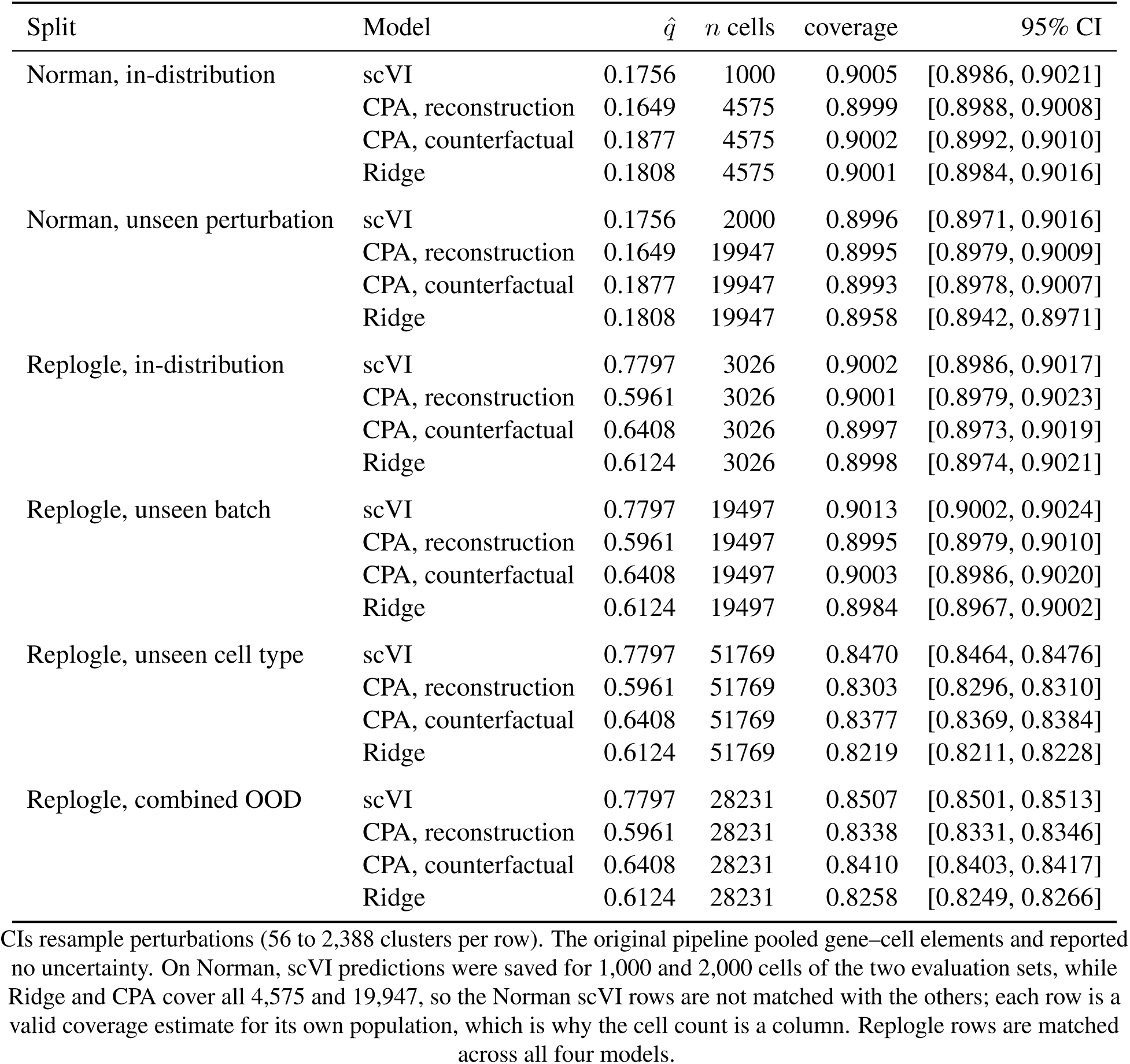
Split-conformal coverage at a nominal 0.90, calibrated on a held-out half of the in-distribution split and scored on disjoint cells. *q*^uses the finite-sample level ⌈(*n*+1)(1−*α*)⌉*/n*. Coverage is the marginal fraction of (cell, gene) pairs inside ±*q*^, *n* counts cells, and the CI is a 95% percentile interval from resampling perturbations with replacement, 2,000 resamples. CPA reconstruction reads the target cell; CPA counterfactual starts from a random control cell of the same split (Table 2).

| Split | Model | $\hat{q}$ | *n* cells | coverage | 95% CI |
|---|---|---|---|---|---|
| Norman, in-distribution | scVI | 0.1756 | 1000 | 0.9005 | [0.8986, 0.9021] |
|  | CPA, reconstruction | 0.1649 | 4575 | 0.8999 | [0.8988, 0.9008] |
|  | CPA, counterfactual | 0.1877 | 4575 | 0.9002 | [0.8992, 0.9010] |
|  | Ridge | 0.1808 | 4575 | 0.9001 | [0.8984, 0.9016] |
| Norman, unseen perturbation | scVI | 0.1756 | 2000 | 0.8996 | [0.8971, 0.9016] |
|  | CPA, reconstruction | 0.1649 | 19947 | 0.8995 | [0.8979, 0.9009] |
|  | CPA, counterfactual | 0.1877 | 19947 | 0.8993 | [0.8978, 0.9007] |
|  | Ridge | 0.1808 | 19947 | 0.8958 | [0.8942, 0.8971] |
| Replogle, in-distribution | scVI | 0.7797 | 3026 | 0.9002 | [0.8986, 0.9017] |
|  | CPA, reconstruction | 0.5961 | 3026 | 0.9001 | [0.8979, 0.9023] |
|  | CPA, counterfactual | 0.6408 | 3026 | 0.8997 | [0.8973, 0.9019] |
|  | Ridge | 0.6124 | 3026 | 0.8998 | [0.8974, 0.9021] |
| Replogle, unseen batch | scVI | 0.7797 | 19497 | 0.9013 | [0.9002, 0.9024] |
|  | CPA, reconstruction | 0.5961 | 19497 | 0.8995 | [0.8979, 0.9010] |
|  | CPA, counterfactual | 0.6408 | 19497 | 0.9003 | [0.8986, 0.9020] |
|  | Ridge | 0.6124 | 19497 | 0.8984 | [0.8967, 0.9002] |
| Replogle, unseen cell type | scVI | 0.7797 | 51769 | 0.8470 | [0.8464, 0.8476] |
|  | CPA, reconstruction | 0.5961 | 51769 | 0.8303 | [0.8296, 0.8310] |
|  | CPA, counterfactual | 0.6408 | 51769 | 0.8377 | [0.8369, 0.8384] |
|  | Ridge | 0.6124 | 51769 | 0.8219 | [0.8211, 0.8228] |
| Replogle, combined OOD | scVI | 0.7797 | 28231 | 0.8507 | [0.8501, 0.8513] |
|  | CPA, reconstruction | 0.5961 | 28231 | 0.8338 | [0.8331, 0.8346] |
|  | CPA, counterfactual | 0.6408 | 28231 | 0.8410 | [0.8403, 0.8417] |
|  | Ridge | 0.6124 | 28231 | 0.8258 | [0.8249, 0.8266] |

CIs resample perturbations (56 to 2,388 clusters per row). The original pipeline pooled gene–cell elements and reported no uncertainty. On Norman, scVI predictions were saved for 1,000 and 2,000 cells of the two evaluation sets, while Ridge and CPA cover all 4,575 and 19,947, so the Norman scVI rows are not matched with the others; each row is a valid coverage estimate for its own population, which is why the cell count is a column. Replogle rows are matched across all four models.

### Reporting per gene set and recalibrating per cell type

Both failures have a standard remedy, and Table 6 measures them instead of recommending them. Calibrating a separate *q*^ on the DE elements and on the rest (Mondrian split conformal^16^) gives each gene set its own target. In distribution, Norman DE-gene coverage rises from 0.342 to 0.898 [0.889, 0.906] for scVI and from 0.271 to 0.901 [0.893, 0.909] for Ridge, while all-gene coverage stays at 0.900. Under unseen perturbation it is 0.889 and 0.874, over only 15 perturbation clusters. The price is width. On Norman the DE-gene interval is 1.42 wide for scVI against 0.34 for the other genes, and 1.85 against 0.35 for Ridge, so an honest DE-gene interval is about four to five times wider than the marginal one suggested. Recalibrating on RPE1, using a random half of the unseen-cell-type split and scoring the other half, restores marginal coverage from 0.847 to 0.900 for scVI and from 0.822 to 0.900 for Ridge. On the combined-OOD cells, which share no cells with either calibration set, it gives 0.903 and 0.902. The two corrections address different failures and are not interchangeable. For scVI’s DE genes on RPE1, coverage is 0.766 with neither, 0.806 with the gene-set split alone, 0.830 with recalibration alone, and 0.901 [0.890, 0.911] with both. Ridge reaches 0.899 on the RPE1 DE genes from the gene-set split alone, but its all-gene coverage stays at 0.821 until it is recalibrated. Gene-set calibration can also over-correct: on the Replogle unseen-batch split, Ridge’s DE intervals cover 0.947. Every coverage here is an empirical fraction over correlated (cell, gene) pairs. The element-level construction does not inherit the finite-sample guarantee, and the bracketed intervals resample perturbations.

**Table 6:**
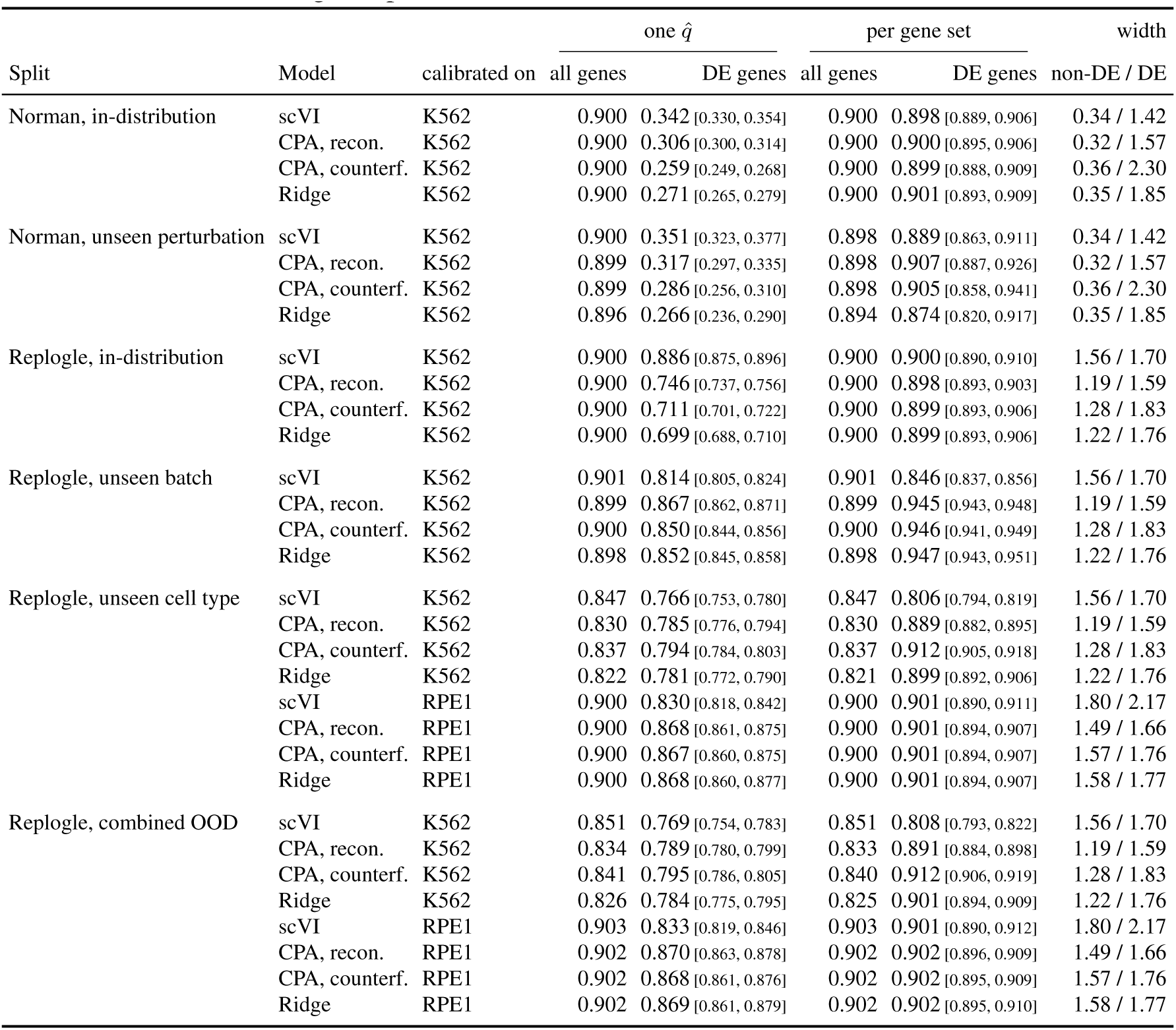
Split-conformal coverage at a nominal 0.90 reported per gene set, and recalibrated when the cell type changes. *One q*^is the construction of Table 5: a single quantile over every calibration (cell, gene) element. *Per gene set* calibrates a separate *q*^ on the DE elements (the 20 DE genes of each cell’s own perturbation) and on the remaining elements (Mondrian split conformal ^16^), so each set has its own target. *Calibrated on* names the cells *q*^was fitted on: K562 is the calibration half of the in-distribution split, RPE1 a random half of the unseen-cell-type split; RPE1 rows on that split are scored on the other half, and K562 rows on the same half, so the two are compared on identical cells. Combined OOD shares no cells with either calibration set. Bracketed intervals resample perturbations with replacement, 2,000 resamples. All coverages are empirical fractions over correlated (cell, gene) pairs.

|  |  |  | one *q̂* |  | per gene set |  | width |
|---|---|---|---|---|---|---|---|
| Split | Model | calibrated on | all genes | DE genes | all genes | DE genes | non-DE / DE |
| Norman, in-distribution | scVI | K562 | 0.900 | 0.342 [0.330, 0.354] | 0.900 | 0.898 [0.889, 0.906] | 0.34 / 1.42 |
|  | CPA, recon. | K562 | 0.900 | 0.306 [0.300, 0.314] | 0.900 | 0.900 [0.895, 0.906] | 0.32 / 1.57 |
|  | CPA, counterf. | K562 | 0.900 | 0.259 [0.249, 0.268] | 0.900 | 0.899 [0.888, 0.909] | 0.36 / 2.30 |
|  | Ridge | K562 | 0.900 | 0.271 [0.265, 0.279] | 0.900 | 0.901 [0.893, 0.909] | 0.35 / 1.85 |
| Norman, unseen perturbation | scVI | K562 | 0.900 | 0.351 [0.323, 0.377] | 0.898 | 0.889 [0.863, 0.911] | 0.34 / 1.42 |
|  | CPA, recon. | K562 | 0.899 | 0.317 [0.297, 0.335] | 0.898 | 0.907 [0.887, 0.926] | 0.32 / 1.57 |
|  | CPA, counterf. | K562 | 0.899 | 0.286 [0.256, 0.310] | 0.898 | 0.905 [0.858, 0.941] | 0.36 / 2.30 |
|  | Ridge | K562 | 0.896 | 0.266 [0.236, 0.290] | 0.894 | 0.874 [0.820, 0.917] | 0.35 / 1.85 |
| Replogle, in-distribution | scVI | K562 | 0.900 | 0.886 [0.875, 0.896] | 0.900 | 0.900 [0.890, 0.910] | 1.56 / 1.70 |
|  | CPA, recon. | K562 | 0.900 | 0.746 [0.737, 0.756] | 0.900 | 0.898 [0.893, 0.903] | 1.19 / 1.59 |
|  | CPA, counterf. | K562 | 0.900 | 0.711 [0.701, 0.722] | 0.900 | 0.899 [0.893, 0.906] | 1.28 / 1.83 |
|  | Ridge | K562 | 0.900 | 0.699 [0.688, 0.710] | 0.900 | 0.899 [0.893, 0.906] | 1.22 / 1.76 |
| Replogle, unseen batch | scVI | K562 | 0.901 | 0.814 [0.805, 0.824] | 0.901 | 0.846 [0.837, 0.856] | 1.56 / 1.70 |
|  | CPA, recon. | K562 | 0.899 | 0.867 [0.862, 0.871] | 0.899 | 0.945 [0.943, 0.948] | 1.19 / 1.59 |
|  | CPA, counterf. | K562 | 0.900 | 0.850 [0.844, 0.856] | 0.900 | 0.946 [0.941, 0.949] | 1.28 / 1.83 |
|  | Ridge | K562 | 0.898 | 0.852 [0.845, 0.858] | 0.898 | 0.947 [0.943, 0.951] | 1.22 / 1.76 |
| Replogle, unseen cell type | scVI | K562 | 0.847 | 0.766 [0.753, 0.780] | 0.847 | 0.806 [0.794, 0.819] | 1.56 / 1.70 |
|  | CPA, recon. | K562 | 0.830 | 0.785 [0.776, 0.794] | 0.830 | 0.889 [0.882, 0.895] | 1.19 / 1.59 |
|  | CPA, counterf. | K562 | 0.837 | 0.794 [0.784, 0.803] | 0.837 | 0.912 [0.905, 0.918] | 1.28 / 1.83 |
|  | Ridge | K562 | 0.822 | 0.781 [0.772, 0.790] | 0.821 | 0.899 [0.892, 0.906] | 1.22 / 1.76 |
|  | scVI | RPE1 | 0.900 | 0.830 [0.818, 0.842] | 0.900 | 0.901 [0.890, 0.911] | 1.80 / 2.17 |
|  | CPA, recon. | RPE1 | 0.900 | 0.868 [0.861, 0.875] | 0.900 | 0.901 [0.894, 0.907] | 1.49 / 1.66 |
|  | CPA, counterf. | RPE1 | 0.900 | 0.867 [0.860, 0.875] | 0.900 | 0.901 [0.894, 0.907] | 1.57 / 1.76 |
|  | Ridge | RPE1 | 0.900 | 0.868 [0.860, 0.877] | 0.900 | 0.901 [0.894, 0.907] | 1.58 / 1.77 |
| Replogle, combined OOD | scVI | K562 | 0.851 | 0.769 [0.754, 0.783] | 0.851 | 0.808 [0.793, 0.822] | 1.56 / 1.70 |
|  | CPA, recon. | K562 | 0.834 | 0.789 [0.780, 0.799] | 0.833 | 0.891 [0.884, 0.898] | 1.19 / 1.59 |
|  | CPA, counterf. | K562 | 0.841 | 0.795 [0.786, 0.805] | 0.840 | 0.912 [0.906, 0.919] | 1.28 / 1.83 |
|  | Ridge | K562 | 0.826 | 0.784 [0.775, 0.795] | 0.825 | 0.901 [0.894, 0.909] | 1.22 / 1.76 |
|  | scVI | RPE1 | 0.903 | 0.833 [0.819, 0.846] | 0.903 | 0.901 [0.890, 0.912] | 1.80 / 2.17 |
|  | CPA, recon. | RPE1 | 0.902 | 0.870 [0.863, 0.878] | 0.902 | 0.902 [0.896, 0.909] | 1.49 / 1.66 |
|  | CPA, counterf. | RPE1 | 0.902 | 0.868 [0.861, 0.876] | 0.902 | 0.902 [0.895, 0.909] | 1.57 / 1.76 |
|  | Ridge | RPE1 | 0.902 | 0.869 [0.861, 0.879] | 0.902 | 0.902 [0.895, 0.910] | 1.58 / 1.77 |

### Decision value: does confidence rank usefully?

Calibration asks whether an interval covers. Ranking asks a different question: given a batch of predictions and budget to validate ten of them, does ordering by the model’s own confidence beat picking ten at random? We rank cells by conformal or quantile interval width and measure the reduction in DE-gene MSE among the top-*k* against random selection, for *k* ∈ {10, 25, 50, 100}, with 95% bootstrap intervals over 2,000 resamples. For scVI both widths are functions of a prediction that the encoder computed from the measured expression of the very cell being ranked, so the scVI rows of Table 7 report the confidence scores a leaky harness produces, not signals that were available before the cell was measured. Residual-based confidence is excluded on principle: it requires the ground truth and amounts to oracle ranking.

**Table 7:** Decision value under convention D: percentage reduction in DE-gene MSE from selecting the *k* most confident predictions, against random selection. *^∗^* marks a comparison whose perturbation-cluster bootstrap *p*-value stays below the Benjamini–Hochberg threshold at *q* = 0.05 across the 48 split × signal × *k* comparisons; the bracketed intervals are 95% percentile intervals from resampling perturbations with replacement. No ground truth enters the ranking; for scVI the ranking signal is a function of a prediction the encoder computed from the ranked cell’s own measured expression. Ties are resolved by the exact expectation over orderings rather than by argsort, which is why no point estimate falls outside its own interval. The in-distribution normalized-conformal rows exclude the calibration half the difficulty model was fitted on, so they and the original-heads rows of the same split are not scored on the same cells. DE-gene MSE is per cell, over the 20 DE genes of that cell’s perturbation, so the unit ranked and selected is the cell and the percentages compare mean per-cell error. The models column lists the models each row covers (S, scVI; R, Ridge). CPA is in no row: its per-cell predictions were regenerated for this revision (Table 2), but the uncertainty methods scored here were not rerun for it.

| Split | Signal | *k*=10 | *k*=25 | *k*=50 | *k*=100 | models |
|---|---|---|---|---|---|---|
| Norman, in-distribution | conformal (normalized) | +2.3 [-47,+52] | +13.1 [-17,+41] | +21.1* [+4,+39] | +24.5* [+15,+34] | S |
|  | quantile (original heads) | +34.3 [-2,+46] | +15.9 [-6,+33] | +19.0* [+4,+30] | +6.8 [-6,+20] | S |
|  | quantile (refit heads) | +52.5 [+37,+62] | +41.6 [+29,+52] | +35.9 [+25,+43] | +27.9 [+21,+35] | S |
| Norman, unseen perturbation | conformal (normalized) | +39.9* [+20,+58] | +43.4* [+30,+57] | +47.3* [+35,+54] | +35.6* [+28,+44] | S |
|  | quantile (original heads) | −3.7 [-48,+37] | −2.0 [-26,+20] | −0.8 [-13,+16] | +2.6 [-10,+18] | S |
|  | quantile (refit heads) | +50.6 [+39,+63] | +48.9 [+38,+57] | +45.0 [+34,+53] | +38.1 [+30,+44] | S |
| Replogle, in-distribution | conformal (normalized) | −4.7 [-44,+43] | −12.9 [-59,+25] | −7.4 [-43,+19] | −0.3 [-20,+21] | S |
|  | quantile (original heads) | −52.6* [-119,-12] | −66.5* [-106,-26] | −59.0* [-90,-28] | −45.9* [-68,-23] | S |
| Replogle, unseen batch | conformal (normalized) | −47.4* [-90,-8] | −21.7* [-56,-4] | −20.7 [-44,-1] | −0.5 [-15,+12] | S |
|  | quantile (original heads) | −30.0 [-78,+0] | −21.3 [-52,-1] | −21.3* [-39,-4] | −24.4* [-38,-11] | S |
| Replogle, unseen cell type | conformal (normalized) | −35.6 [-105,+8] | −26.7 [-64,+1] | −39.8* [-65,-10] | −31.9* [-54,-18] | S |
|  | quantile (original heads) | −44.5* [-81,-21] | −55.3* [-82,-31] | −46.3* [-70,-26] | −40.4* [-57,-25] | S |
| Replogle, combined OOD | conformal (normalized) | −16.4 [-63,+12] | −27.2* [-57,-5] | −42.2* [-62,-19] | −42.7* [-61,-24] | S |
|  | quantile (original heads) | −65.5* [-138,-21] | −53.8* [-90,-23] | −43.2* [-71,-26] | −54.2* [-73,-36] | S |

2,000 resamples. Of the 48 comparisons, 28 remain significant after correction with perturbation-level resampling, against 30 with cell-level resampling and 31 with perturbation-within-batch resampling on Replogle. The refit-heads rows are outside that 48-comparison family and carry no asterisk; each of their perturbation-level intervals excludes zero, and in distribution they rank the 909 DE-evaluable cells of the evaluation half rather than every cell. Ridge is omitted: its Norman unseen-perturbation prediction has zero across-cell variance, so every label-free signal derived from it is constant and there is no ranking to evaluate. Table 9 gives the distinct-value count for every Ridge signal on every split.

Table 7 reports the result, and the pattern is a property of what the model was given rather than of the ranking signal. On Norman, where scVI is conditioned on guide_id, conformal width cuts DE-gene error by 35.6–47.3% against random at every *k* on the unseen-perturbation split, with all four intervals excluding zero. On Replogle, where the checkpoint was trained with no perturbation covariate at all (“Conditioning covariates absent at training time”), the same signal is negative on every row and significantly so on seven of sixteen after correction. A confidence signal cannot rank by a variable the model was never given; it inherits the conditioning of the model that produced it, so the two datasets cannot be pooled into a single verdict.

The two Ridge rows are omitted from Table 7 rather than reported as numeric decision values, because the Norman unseen-perturbation prediction has zero across-cell variance and every label-free signal derived from it is constant (Table 9, Supporting Online Material). Ridge and, since this revision, CPA’s counterfactual arm are the only genuine counterfactual predictions, since neither receives the test cell’s expression; on Norman unseen perturbation Ridge’s prediction is constant across cells, so its ranking is undefined rather than poor, and no counterfactual ranking result exists for scVI, which has no counterfactual arm, or for CPA, for whose counterfactual no confidence signal was computed.

### Calibration and ranking are different properties

Table 4 places the three UQ methods side by side on scVI, Norman, unseen perturbation, at a nominal 0.90. MC dropout is the *worst*-calibrated method by a wide margin (0.0191 empirical coverage against a nominal 0.90) and simultaneously the *best* ranking signal, at +63.6% [+45.7, +70.7] on the top ten; the gap is widest at small *k* (Figure 4, Supporting Online Material). Plain conformal is nearly exactly calibrated (0.8996) but has a single width and cannot rank; the normalized conformal interval, which can, covers 0.4535 of cells and ranks worst of the three ranking signals (+39.9%). Ordered by calibration error the three ranking signals run refit quantile, normalized conformal, MC dropout; ordered by top-10 gain they run MC dropout, refit quantile, normalized conformal (Figure 2). A benchmark that reports only coverage would select plain conformal, which cannot rank at all, and discard the signal that ranks best; one that reports only top-*k* gain would select intervals covering 1.9% of the time at a nominal 90%. Both numbers are needed, and reporting one is not evidence about the other^26^.

**Figure 2:**
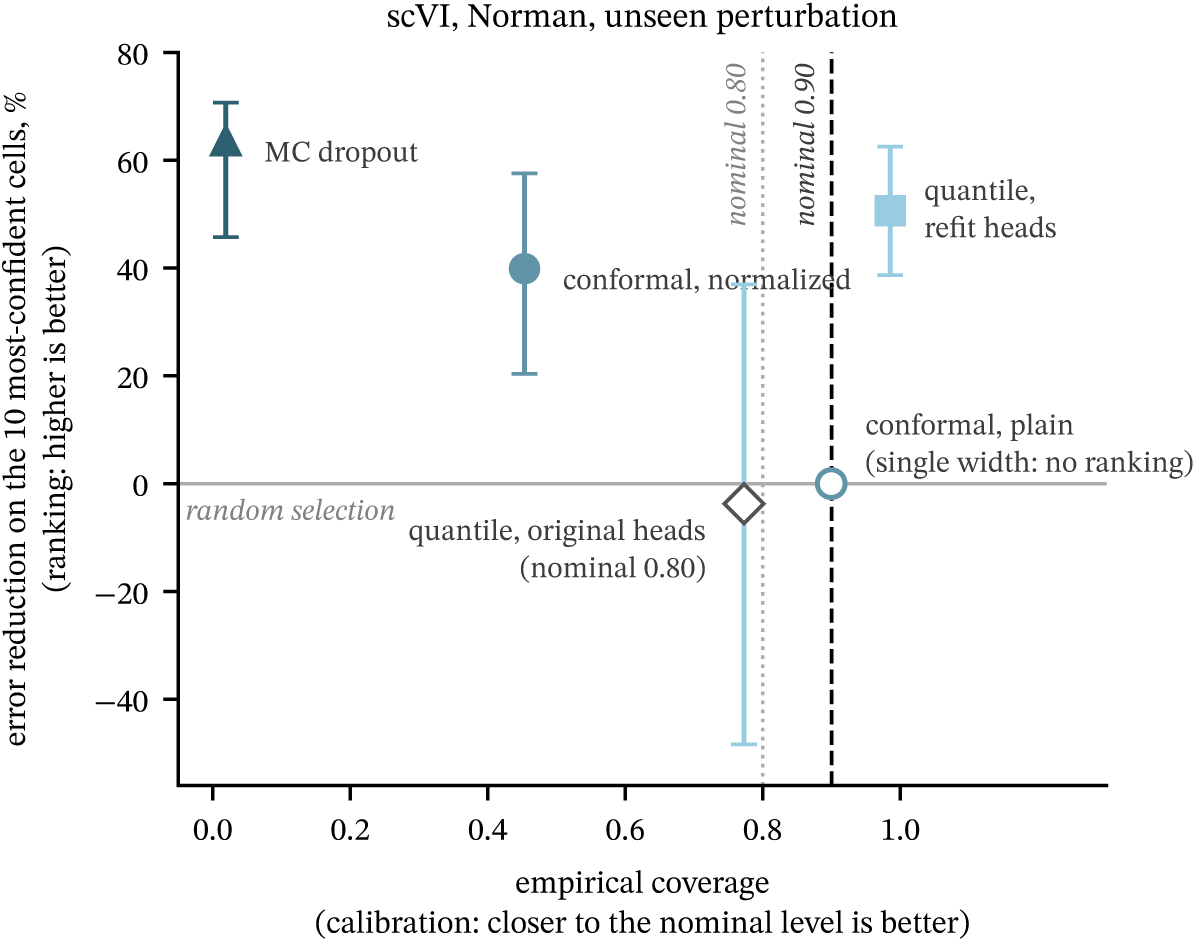
Calibration and ranking are different properties. MC dropout is the worst-calibrated of the five intervals and simultaneously the best ranking signal. Each row of Table 4 is one point: plain conformal is drawn hollow on the random-selection line because its single width defines no ranking; the normalized conformal point sits at the coverage of its own interval (0.4535); the refit-heads point is the replicated gain (+50.6); and the original heads sit at their stored coverage at nominal 0.80. Bars are 95% bootstrap intervals over the 15 perturbations. Plotted with matplotlib. Coverage is over (cell, gene) pairs; the error reduction is over per-cell DE-gene MSE.

## Discussion

### Failure modes

Assessing whether perturbation models are reliable in practice takes more than measuring their accuracy; measuring it the wrong way is worse than not measuring it, because a harness that is wrong in these ways still reports success^28^. Each failure mode below has that signature. The configuration is self-consistent, the code executes as written, and every reported number falls in a plausible range, so nothing in the output separates the broken measurement from a sound one; what separates them is a null baseline the harness never ran. Each is given below as a mechanism, the diagnostic that reveals it, and a detection method transferable to other pipelines.

### Assessing the reconstruction as though it were a prediction

The harness ran each model’s inference routine on the unaltered test object and it compared the output against that object’s expression matrix. The encoder saw the target cell before the decoder generated the profile it was scored against. This behaviour is documented and is not a coding error: CPA’s predict() exposes both a reconstruction and a counterfactual mode, and returns the reconstruction by default. The default is not a property of our configuration. predict() exposes both modes and returns the reconstruction unless a counterfactual is requested, so any harness that calls it without naming the mode scores reconstruction fidelity under a perturbation-effect label, and an identity baseline is the only thing that separates the two.

As a diagnostic, report the score of an identity predictor that returns each cell’s own expression. With the effect-correlation approach employed here, the predictor achieves an effect correlation of 1.000 on all six splits when returned unconverted, and 0.997 to 1.000 under convention D (Table 2), meaning that any model that is given the target cell is being assessed against an echo ceiling instead of a null.

For detection, verify the model’s input at inference time. When the input tensor includes the target cell’s expression, the metric measures reconstruction fidelity rather than prediction^5^.

### Metrics a constant predictor already saturates

Correlating mean profiles at the dataset level is insensitive to perturbation identity, since averaging occurs before correlation is computed. On Norman in-distribution, a constant control-mean predictor scores *r* = 0.9983 under this definition and the Ridge baseline scores 0.99995, against 1.0000 for an identity predictor that simply returns its input: the entire scale between carrying no perturbation information and perfect reconstruction is 0.0017 wide. The problem is worse than a saturated metric, because the original study’s point-accuracy table did not use a single definition throughout: its scVI and Ridge values were element-wise correlations over all (cell, gene) pairs, while its CPA value of 0.9995 was a profile correlation between mean expression vectors. Two rows of the same column reported different statistics. Table 2 reports the element-wise statistic for every row and states so in its caption. Two independent benchmarks explain why: Perturb-seq benchmarks carry low perturbation-specific variance, so a training-set-mean baseline beats scGPT and scFoundation^12^, and a perturbation-informed mean control exceeds GEARS and scGPT^11^; a mean-based predictor is a strong null, which is what makes a metric it saturates uninformative.

The same insensitivity appears in the DE-gene column: under the published convention (A), in which the original table was computed, a predictor that outputs zero for every gene scores a DE-gene MSE of 3.849 on Norman in-distribution against the 1.652 we reported for scVI. Table 2 reports convention D, under which those scVI predictions score 0.1921; the zero predictor scores 3.849 under either, since rescaling a zero prediction leaves it zero.

As a diagnostic, evaluate the null predictor using the same metric definition that was used for the models, not by reimplementing that definition, and confirm that every row of a reported column uses the same definition as every other.

For detection, if the null value is within a few thousandths of the best model, then the metric will not be able to support a claim regarding what the model has learned.

### Holding out the wrong column

In the Replogle split configuration the column held out was designated as batch, that is, an integer batch identifier, rather than as perturbation identity. The 25% holdout was applied accurately and correctly, 12 out of the 48 K562 batches, but was done with respect to a variable that was not the one intended for the generalisation test. Of the 2,007 perturbation labels in the resulting ‘unseen-perturbation’ split, 2,004 appear in the training set and 0.03% of the cells have a perturbation which was not present in the training data. This split assesses reconstruction in the case of batch shift. The corresponding Norman configuration, on the other hand, named the right column and has a genuine holdout, even though this is 20% rather than the 25% originally reported. We therefore relabel the Replogle split *unseen batch* in every table and figure, and rebuild the split on the intended column (“Methods”): on the rebuilt split, Ridge conditioned on gene carries perturbation signal in distribution (effect *r* = 0.197, against 0.002 under batch conditioning), and on the 410 held-out genes every predictor that is denied the target cell is matched or beaten by the train-mean floor (Table 3).

As a diagnostic, present the |test labels \ train labels| and the fraction of test cells that have a novel label, for each split, in the form of a table row rather than as a configuration value^24^.

For detection, the fact that a holdout achieves exactly its nominal rate is proof that the code was executed, not evidence that the intended variable was held out.

### Conditioning covariates absent at training time

The Replogle scVI checkpoint carries no additional categorical covariates and a batch key with a single category: it was given no perturbation identity at training or at inference, so every Replogle scVI entry in our results refers to an autoencoder that cannot distinguish perturbations. It nevertheless beats a constant on the DE genes in distribution, because condition-specific deviation enters through the encoder reading each test cell’s expression. An unintentional ablation makes the same point for CPA. Four checkpoints trained against a mismatched perturbation vocabulary hold an identically zero embedding matrix and route none of the 1,743 test perturbations, against 1,742 of the 1,743 for the correctly configured checkpoints. Both arms sit below the control-mean floor of 0.4079 by a similar margin, and the zero-embedding arm scores marginally better on DE-gene MSE (0.3142 against 0.3207, over 4,615 cells) with a higher across-condition variance ratio (0.132 against 0.119). The difference is about 2% of the score and we do not claim the ablation wins; the claim is that the perturbation pathway confers no measurable advantage, and that whatever advantage either arm holds over the floor comes from the encoder reading the test cell rather than from perturbation identity.

As a diagnostic, look at the saved model registry, not the training script, since registries keep a record of what the model was actually constructed with.

For detection, zero the perturbation input and re-score. If the metric is unchanged, the pathway is not contributing.

### Silent substitution of unseen categories

Models which do not have unseen perturbation labels need a fallback option, and our harness included seven different ones. Three illustrate the range. When dealing with unseen Norman guides, the system refers to an existing negative-control guide; the conformal method resolves these cases to a different sentinel because the intended negative control is not present in the training vocabulary and as a result the code defaults to the label that comes first in alphabetical order; the Ridge baseline is given a feature row consisting of all zeros and then returns its intercept, so that its predictions on that split are constant across cells to within 1 × 10^−6^. In the training vocabulary of the stored split the negative-control rule (training notebook cells 177 and 195) resolves to AHR_NegCtrl0;AHR_NegCtrl0, a single-gene perturbation paired with a control guide, and the alphabetical rule (cells 159, 161 and 163) to AHR_FEV;AHR_FEV. All of the entries in the Norman unseen-perturbation split are impacted. Two of these three are found in the same notebook and serve the same purpose, yet resolve to different labels. On Replogle the corresponding rule (training notebook cell 273) finds no training label containing ctrl or neg and resolves to the alphabetically first label, the gene AAAS.

As a diagnostic, break down the number of cells that received a substituted label and state what each substitution amounted to in the training vocabulary, not what the source code names were.

For detection, a sentinel string which is not included in the vocabulary will not trigger; instead it will quietly take on the value chosen by the fallback.

### Conclusions that depend on an arbitrary convention

When predictions and targets are given in different units, a conversion is necessary and several reasonable conventions exist^29^. For the Replogle in-distribution row that carries the largest decisionvalue move, conventions B and D change all-gene MSE by 0.01%; across the full harness (Figure 5, Supporting Online Material) the largest change is 4.7% of the convention-B value and 8 of the 12 model-split rows move by more than 1%. The convention choice nevertheless moves the most convention-sensitive decision-value entry by 29.1 percentage points and reverses 5 of the 48 significance judgments in Table 7; including the Ridge rows in Table 9 adds 4 more, all from a single ranking signal that takes 36 distinct values across 4,615 cells. That entry is Replogle in-distribution, conformal width at *k*=10, which reads −32.7% under the asymmetric convention and −3.5% under D. A different entry on the same row, at *k*=100, moves from +23.8% [+7.8, +36.6] to +2.4% [−16, +20] and loses significance, under a convention chosen on principle rather than to obtain that outcome.

As a diagnostic, recompute all the headlines under each defensible convention and state the spread, not just the preferred value.

For detection, a conclusion that moves more under a unit convention than under a change of model is a conclusion about the convention^28^.

## Conclusion

CPA’s regenerated predictions enter the point-accuracy and coverage tables; the comparisons of uncertainty methods below rest on scVI and Ridge, and CPA’s negative-binomial-on-log1p configuration is reported as trained, not tuned. The harness has two genuinely counterfactual predictors. The linear baseline matches its train-mean floor on Norman unseen perturbation (Table 8, Supporting Online Material), and CPA’s counterfactual arm has a higher error than a constant profile on every split (Tables 2 and 11); it beats Ridge only on the two RPE1 splits, where its input cells carry the cell line that Ridge never sees. The question Ahlmann-Eltze et al.^2^ answered in the negative on data of this kind, whether deep perturbation models outperform simple linear baselines, can therefore be posed for CPA, and on these outputs it does not, for one seed and one configuration that applied a count likelihood to log1p input. It still cannot be posed for scVI, which has no counterfactual arm. In the configurations tested, split conformal prediction was the only method to attain its nominal coverage in distribution, and it lost 0.053 of that under cell-type shift on Replogle, so exchangeability would have to be re-established by recalibrating on the target cell type^27^. Quantile regression as originally configured undershot at every nominal level, and a monotonicity constraint removes the crossing but leaves scVI over-covering (0.913–0.985). On scVI, Norman, unseen perturbation, MC dropout is the worst-calibrated of the three methods and simultaneously the best ranking signal: calibration and ranking are different properties, and a benchmark reporting only the first would order these three implementations backwards. Confidence-ranked selection has no single verdict across the two datasets, because a confidence signal inherits the conditioning of the trained configuration that produced it. On Norman, where scVI is conditioned on guide_id, conformal width cuts DE-gene error against random selection at every *k*; on Replogle, where the checkpoint received no perturbation covariate at all, the same signal is negative on almost every row.

These findings support split conformal prediction for marginal all-gene coverage on the two datasets and the three models for which per-cell predictions now exist (CPA’s coverage behaves as scVI’s and Ridge’s do, Tables 5 and 6), with recalibration whenever the target cellular context changes. They do not support the same single-quantile intervals for DE-gene uncertainty: even in distribution, DE-gene coverage is far below nominal. Calibrating per gene set and recalibrating per cell type restore it (Table 6), at the cost of DE-gene intervals four to five times wider than the marginal one, and both corrections are needed under cell-type shift. A benchmark should therefore report coverage per gene set, and recalibrate whenever the cellular context changes. These are results on two datasets, three models with per-cell predictions and one seed, and they need replication before they are read as properties of the methods. Future work will include adding more models, scaling CPA to larger perturbations, using more datasets (e.g. Adamson 2016^1^) and saving per-cell predictions across all models.

### Limitations

There are limits to our audit worth making clear. Only Ridge and CPA’s regenerated counterfactual arm carry out actual counterfactual prediction; scVI has no counterfactual arm. The audit covers two datasets and three models, and CPA values come from a single seed. Norman’s unseen-perturbation holdout is 20% of guide identities rather than the 25% first described, and after negative-control substitution every held-out guide reaches the model as the control label, so no model is assessed on genuinely novel perturbations in that split. Some of these are findings rather than caveats: a split that silently substitutes every held-out label is a flaw in the evaluation design, not a limitation of scope. A counterfactual rerun of the decision-value analysis, in which scVI and CPA would receive only control-cell expression, the perturbation identity and covariates known before measurement, would require retraining both models with the perturbation covariate as an input, since the Replogle scVI checkpoint received none and the CPA checkpoints could not use it, and was not performed.

Two kinds of result in this paper should be kept apart. The diagnostics transfer to other harnesses without replication: scoring an identity predictor that returns each cell’s own expression, tabulating the held-out variable and the fraction of test cells with a novel label for every split, and zeroing a model’s perturbation input and re-scoring. The quantitative results are specific to the trained configurations audited here, two datasets, scVI, CPA and Ridge, with CPA from one seed and without uncertainty methods, and one implementation of each uncertainty method, and they need replication on other datasets, models and implementations before they can be read as properties of the architectures. In particular, the finding that the Replogle scVI checkpoint and the CPA checkpoints could not use perturbation identity describes how those models were configured and trained in this pipeline, not what the architectures can do.

## Acknowledgments

This work was conducted through the Algoverse AI Research program. We thank our mentor and the program staff for guidance.

## AI use statement

We used LLM-based coding assistants (Claude via Claude Code) during the audit reported here. Their role was mechanical: re-deriving metrics from the stored per-cell prediction arrays, and verifying tables against our scripts’ CSV output. Every number reported here was produced by released code and checked against the underlying arrays by the authors. All prose in this manuscript was written by the authors; no claim, experimental design, interpretation, or sentence was generated or rewritten by a language model.

