## Supporting Online Material for "Auditing Confidence-Calibrated Benchmarking of Single-Cell Perturbation Models"

**Manuel Gallone***, **Shyam Ravidath***, **Srihith Madireddy**
Algoverse AI Research


### Supporting Online Material

The tables and figures in this appendix support "Results". They are reported under the same constant-library convention (D) as the main text.

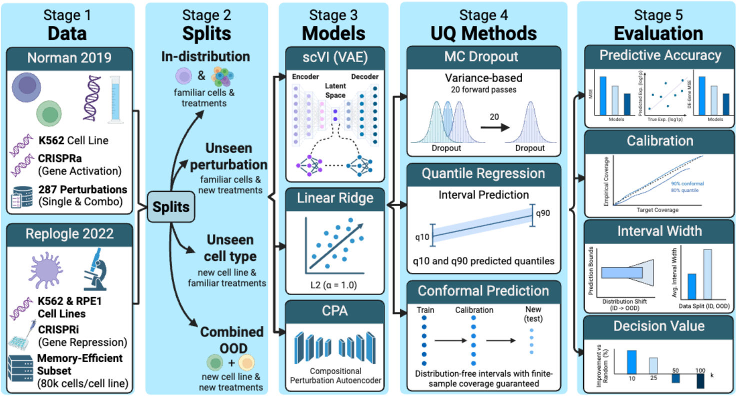


**Figure 3:** Five-stage pipeline for uncertainty-aware OOD evaluation of perturbation models. Stage 1 ingests two datasets (Norman 2019, CRISPRa K562; Replogle 2022, CRISPRi K562+RPE1), and Stage 2 partitions them into four splits: in-distribution, unseen perturbation, unseen cell type, and combined OOD. Stage 3 trains three models of varying complexity (scVI VAE, Linear Ridge, CPA), and Stage 4 applies three UQ methods (MC Dropout, quantile regression, split conformal prediction) to each. Stage 5 evaluates results across five axes: predictive accuracy (MSE, Pearson $r$, DE-gene MSE), calibration, interval width, and decision value. Created with BioRender.

*Equal contribution (co-first authors).

**Table 6:** Every split, and the column it was built on. The Replogle split configured as unseen perturbation, here *unseen batch*, was built on `batch` rather than on perturbation identity; "Holding out the wrong column" reports what that does to the split. The held-out variable is none for in-distribution, `guide_id` for Norman unseen perturbation, `batch` for Replogle unseen batch, `cell_type` for unseen cell type and `batch` together with `cell_type` for combined OOD; the four right-hand columns count unique values of the `perturbation` label in the stored split files: perturbations in the training set, in the test set, in the test set but absent from training, and the percentage of test cells carrying such an absent label.

| Dataset | Split | Held-out column | Holdout | perts (train) | perts (test) | absent | % cells novel |
|---|---|---|---|---|---|---|---|
| Norman 2019 | in-distribution | none | test 10% of K562 | 197 | 197 | 0 | 0.0 |
| CRISPRa, K562 | unseen perturbation | `guide_id` | 20% of guide identities | 197 | 56 | 40 | 61.4 |
| Replogle 2022 | in-distribution | none | test 10% of K562 | 2050 | 1743 | 1 | 0.02 |
| CRISPRi, | unseen batch | `batch` | 12 of 48 K562 batches | 2050 | 2007 | 3 | 0.03 |
| K562 and RPE1 | unseen cell type | `cell_type` | all RPE1 cells | 2050 | 2388 | 345 | 15.3 |
| | combined OOD | both | RPE1 cells with held-out batches | 2050 | 2331 | 335 | 15.5 |

Norman carries 290 guide conditions (237 perturbation labels) and 33,694 genes before filtering, of which 5,000 highly variable genes are kept; train, calibration and in-distribution test are 80/10/10 of K562. Replogle uses 80,000 cells per cell line under the same 80/10/10 on K562. No model predictions were retained over the Replogle calibration cells, which is why $\hat{q}$ is calibrated on a split half of the in-distribution test set ("Uncertainty Quantification and Evaluation").

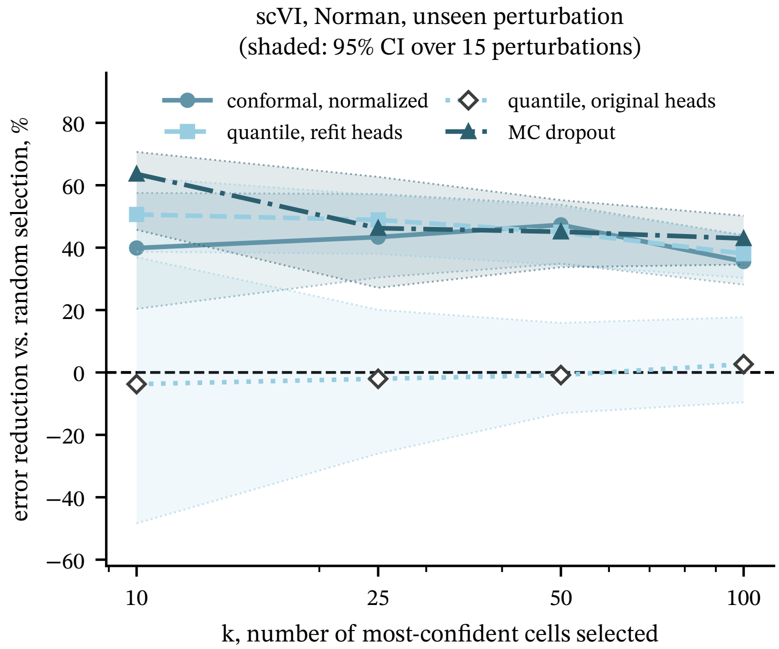


**Figure 4:** Decision value against the number of selected cells $k$ for the four ranking signals of Table 4, with 95% bootstrap intervals over the 15 perturbations; plain conformal has a single width and no curve. The advantage of MC dropout is largest at small $k$, which is the regime a prioritization budget actually operates in. Plotted with matplotlib. The error reduction is computed over per-cell DE-gene MSE.

**Table 7:** Training configuration for every model and uncertainty method. Reported for reproducibility; no claim in this paper turns on any of these settings. CPA ran under Python 3.10 with cpa-tools 0.8.8, scvi-tools 0.20.3, PyTorch 2.0.0 and anndata 0.9.2. The refit quantile heads added in revision ("Uncertainty Quantification and Evaluation") are listed separately from the original heads.

| Component | Aspect | Value |
|---|---|---|
| scVI | architecture | `n_latent=64` (Norman), 128 (Replogle); `n_layers=2`; `dropout_rate=0.1` |
| | optimisation | Adam, lr 3e-4, `max_epochs=200`, early stopping patience 15 |
| | prediction | `get_normalized_expression()`, proportion space (0–0.004) |
| | seeds | 42, 0, 1, 7, 123 |
| CPA | architecture | `n_latent=128`, `recon_loss="nb"`, `doser_type="linear"` |
| | optimisation | `max_epochs=80`, `batch_size=64`, early stopping patience 5 |
| | perturbation key | `guide_id` on Norman, every guide pair containing a NegCtrl guide (3 control and 125 single-gene pairs) collapsed to a single control label |
| | seeds | 42, 0, 1, 7, 123 |
| Ridge | features | one-hot: perturbation identity (Norman), sequencing batch (Replogle); no cell-type covariate |
| | regularisation | $\alpha = 1.0$; 5-member ensemble over $\alpha \in \{0.5, 1.0, 2.0, 5.0, 10.0\}$, inter-member spread as the uncertainty signal |
| | fitting | deterministic, trained once |
| MC dropout | passes | 20 forward passes for scVI, 5 for CPA |
| Quantile regression, original heads | heads | three linear heads on scVI's latent space at the 10th, 50th and 90th percentile, applied unchanged to the Ridge baseline |
| | training | 30 epochs, Adam, lr 1e-3, pinball loss |
| | nominal levels | 0.80, 0.90, 0.95 |
| Quantile regression, refit heads | features | top-10 principal components of scVI's predicted profile under convention D, fitted on the fitting half of the matched in-distribution cells |
| | heads | three linear heads at the 5th, 50th and 95th percentile (nominal 0.90), made monotone by sorting the three predicted surfaces elementwise at prediction time (rearrangement) |
| | training | 300 Adam steps, pinball loss, warm start at the empirical per-gene quantiles, step size $0.05\times$ the warm-start interval width with cosine decay; pinball loss 0.0124 at the warm start, 0.0116 at the end |
| | result | crossing rate 0.0; element coverage 0.982 in distribution, 0.985 unseen perturbation |
| Split conformal | level | $\alpha = 0.10$, target coverage 0.90 |
| Decision value | budgets | $k \in \{10, 25, 50, 100\}$ |
| | intervals | 95% bootstrap over 2,000 resamples |

**Table 8:** Each model against a constant train-mean predictor that ignores its input. *MSE / floor* is the ratio of mean per-cell squared error; *win rate* is the fraction of cells on which the model beats the constant (0.5 is a coin flip); the CI is a paired bootstrap over cells on the per-cell difference. Win rate and verdict can disagree, and the two Ridge rows on Replogle are the clearest case: at a win rate of 0.372 on unseen cell type the model loses on most cells, yet wins by enough on the rest that the mean paired difference is negative and its interval excludes zero. The win rate counts cells; the verdict is the sign of the mean. A skewed per-cell difference distribution separates the two, and "the average improves while most cells get worse" is itself a measurement-validity result rather than a transcription error. The models column lists the models each row covers (S, scVI; R, Ridge). CPA's regenerated predictions are scored against the same floors in Table 2, and its counterfactual arm against the mean of its own input control cells in Table 11.

| Split | Model | MSE / floor | win rate | 95% CI on $\Delta$ | verdict | models |
|---|---|---|---|---|---|---|
| Norman, in-distribution | Ridge | 0.9326 | 0.782 | [-2.19e-03, -1.81e-03] | **beats floor** | S,R |
| | scVI | 0.7615 | 0.997 | [-7.41e-03, -6.76e-03] | **beats floor** | S,R |
| Norman, unseen perturbation | Ridge | 0.9996 | 0.403 | [-3.56e-05, +1.19e-05] | **= floor** | S,R |
| | scVI | 0.7914 | 0.993 | [-6.43e-03, -5.85e-03] | **beats floor** | S,R |
| Replogle, in-distribution | Ridge | 0.9903 | 0.675 | [-1.49e-03, -1.36e-03] | **beats floor** | S,R |
| | scVI | 1.6389 | 0.002 | [+9.35e-02, +9.44e-02] | *worse than floor* | S,R |
| Replogle, unseen batch | Ridge | 1.0001 | 0.371 | [+1.30e-05, +1.53e-05] | *worse than floor* | S,R |
| | scVI | 1.6220 | 0.003 | [+9.15e-02, +9.20e-02] | *worse than floor* | S,R |
| Replogle, unseen cell type | Ridge | 0.9984 | 0.372 | [-4.60e-04, -3.60e-04] | **beats floor** | S,R |
| | scVI | 1.2851 | 0.004 | [+7.18e-02, +7.21e-02] | *worse than floor* | S,R |
| Replogle, combined OOD | Ridge | 0.9996 | 0.748 | [-1.02e-04, -9.90e-05] | **beats floor** | S,R |
| | scVI | 1.2862 | 0.006 | [+7.08e-02, +7.12e-02] | *worse than floor* | S,R |

Paired over cells, 2,000 resamples. With tens of thousands of cells the test resolves differences of order 0.03%, so "= floor" is a strong statement and a significant verdict need not mean a large one.

**Table 9:** Every label-free ranking signal derived from the Ridge baseline is degenerate. *distinct* counts the values the signal takes across the evaluable cells of that split. The counts recover the split configuration rather than any property of the predictions: 36 is the number of sequencing batches present in training, of 48 in the K562 pool; 12 is the number held out; 13 is those same 12 plus one shared value for the 8 batch labels that appear only in the combined-OOD split and are therefore absent from the training vocabulary, which fall back to the intercept; and 1 means the predictor is constant across every cell. Top-$k$ selection over such a signal is decided by tie-breaking, so no improvement percentage is reported. The models column lists the model each row covers (R, Ridge). CPA is in no row: no confidence signal was computed for it.

| Split | Signal | distinct | cells | what the count is | models |
|---|---|---|---|---|---|
| Norman, in-distribution | conformal width | **229** | 8,276 | training perturbations | R |
| | quantile width | **229** | 8,276 | training perturbations | R |
| Norman, unseen perturbation | conformal width | **1** | 4,747 | predictor is constant | R |
| | quantile width | **1** | 4,747 | predictor is constant | R |
| Replogle, in-distribution | conformal width | **36** | 4,615 | sequencing batches | R |
| | quantile width | **36** | 4,615 | sequencing batches | R |
| Replogle, unseen batch | conformal width | **1** | 11,807 | predictor is constant | R |
| | quantile width | **12** | 11,807 | held-out batches | R |
| Replogle, unseen cell type | conformal width | **36** | 21,040 | sequencing batches | R |
| | quantile width | **36** | 21,040 | sequencing batches | R |
| Replogle, combined OOD | conformal width | **1** | 11,482 | predictor is constant | R |
| | quantile width | **13** | 11,482 | held-out batches | R |

Convention D. The largest count anywhere in this table is 229 distinct values over 8,276 cells; the smallest is 1 over 11,807.

**Table 10:** The rebuilt Replogle K562 split, held out on `gene`. Built from the same 80,000 K562 cells as the stored splits; 20% of the 2,050 gene labels (410, seed 20261120) are held out, stratified on whether the gene carries a DE list. *Novel* counts labels absent from the training split, and the percentage is over perturbed (non-control) cells. Compare the stored unseen-batch split (Table 6): 3 novel labels and 0.03% of cells.

| Split | cells | controls | batches | labels | novel | % novel | with DE list |
|---|---|---|---|---|---|---|---|
| training | 51,093 | 1,899 | 48 | 1,643 | 0 | 0.0 | 733 |
| calibration | 6,482 | 280 | 48 | 1,480 | 0 | 0.0 | 716 |
| in-distribution test | 6,318 | 264 | 48 | 1,458 | 0 | 0.0 | 711 |
| gene-holdout test | 16,107 | 288 | 48 | 410 | 410 | 100.0 | 183 |

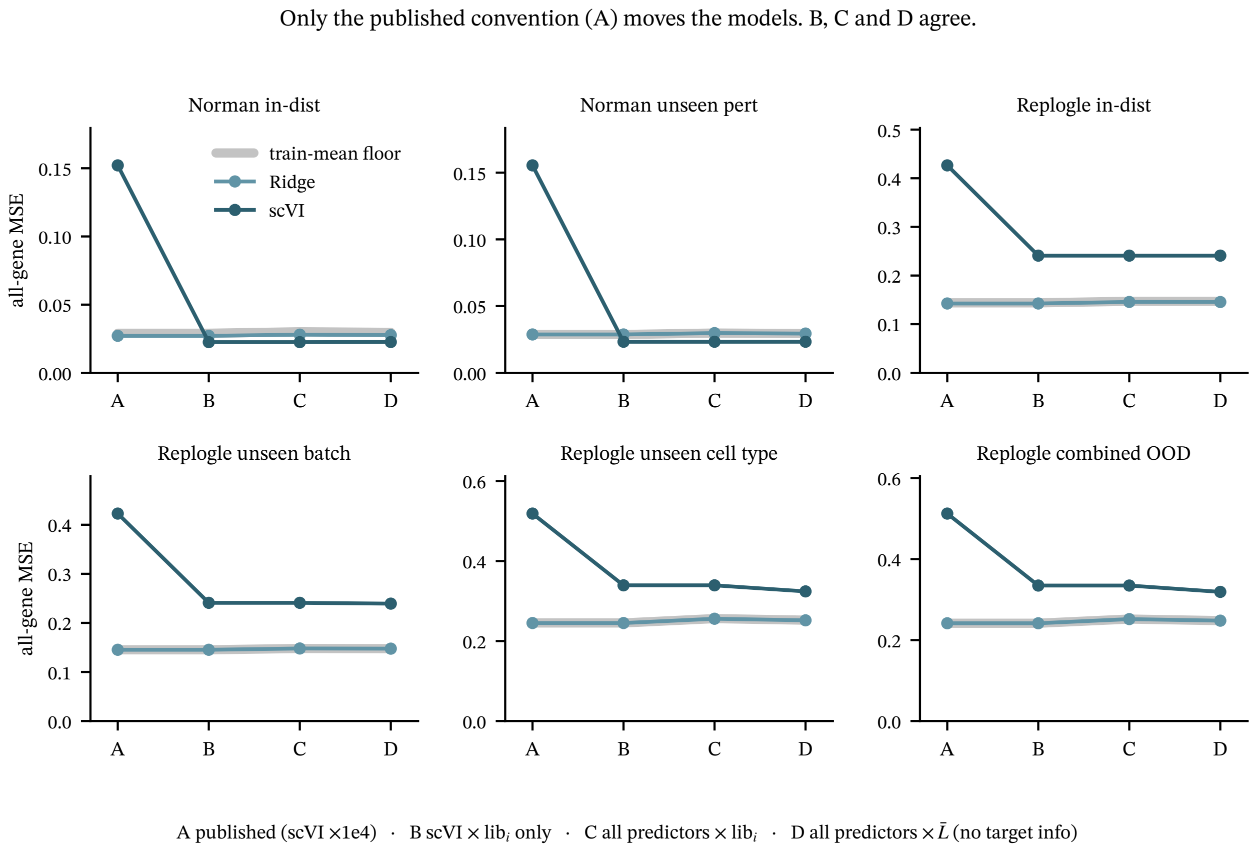


**Figure 5:** All-gene MSE for each predictor under the four library-size conventions, on every split. Only the published convention (A) separates the models; B, C and D agree on the ordering everywhere. Convention D, used throughout this paper, gives no predictor any test-target information. Every panel is anchored at zero, so the drop from A is comparable across panels. The Ridge line rides on the train-mean floor band on five of six splits. Plotted with matplotlib. All-gene MSE is the mean over all (cell, gene) elements of each split.

**Table 11:** The null the CPA counterfactual has to beat. Its input is a control cell drawn at random from the same split, with the target cell's perturbation label, so it sees the split's control state, which on the two RPE1 splits is RPE1. The matching null is a constant: the mean of that same control pool. Convention D, on the cells of Table 2; the DE-gene CI is a bootstrap over cells. On every split, one of the two constants has both a lower all-gene and a lower DE-gene MSE than the counterfactual.

| Split | control pool | Predictor | all-gene MSE | DE-gene MSE | 95% CI |
|---|---|---|---|---|---|
| Norman, in-distribution | 873 | CPA, counterfactual | 0.0349 | 0.5332 | [0.5106, 0.5559] |
| | | *mean of the split's control cells* | 0.0303 | 0.5594 | [0.5331, 0.5842] |
| | | *train-mean floor* | 0.0297 | 0.4964 | [0.4738, 0.5173] |
| Norman, unseen perturbation | 2,948 | CPA, counterfactual | 0.0354 | 0.4669 | [0.4254, 0.5098] |
| | | *mean of the split's control cells* | 0.0302 | 0.4370 | [0.3961, 0.4799] |
| | | *train-mean floor* | 0.0294 | 0.4107 | [0.3734, 0.4493] |
| Replogle, in-distribution | 198 | CPA, counterfactual | 0.1582 | 0.3611 | [0.3543, 0.3674] |
| | | *mean of the split's control cells* | 0.1484 | 0.3606 | [0.3543, 0.3670] |
| | | *train-mean floor* | 0.1470 | 0.3386 | [0.3328, 0.3446] |
| Replogle, unseen batch | 675 | CPA, counterfactual | 0.1585 | 0.2261 | [0.2230, 0.2291] |
| | | *mean of the split's control cells* | 0.1482 | 0.2098 | [0.2069, 0.2126] |
| | | *train-mean floor* | 0.1475 | 0.2065 | [0.2038, 0.2093] |
| Replogle, unseen cell type | 10,369 | CPA, counterfactual | 0.2444 | 0.3133 | [0.3111, 0.3158] |
| | | *mean of the split's control cells* | 0.1644 | 0.2011 | [0.1994, 0.2030] |
| | | *train-mean floor* | 0.2522 | 0.3333 | [0.3308, 0.3365] |
| Replogle, combined OOD | 5,651 | CPA, counterfactual | 0.2410 | 0.3117 | [0.3083, 0.3152] |
| | | *mean of the split's control cells* | 0.1602 | 0.1993 | [0.1967, 0.2018] |
| | | *train-mean floor* | 0.2481 | 0.3301 | [0.3261, 0.3340] |
